# A surface morphometrics toolkit to quantify organellar membrane ultrastructure using cryo-electron tomography

**DOI:** 10.1101/2022.01.23.477440

**Authors:** Benjamin A. Barad, Michaela Medina, Daniel Fuentes, R. Luke Wiseman, Danielle A Grotjahn

## Abstract

Cellular cryo-electron tomography (cryo-ET) enables 3-dimensional reconstructions of organelles in their native cellular environment at subnanometer resolution. However, quantifying ultrastructural features of pleomorphic organelles in three dimensions is challenging, as is defining the significance of observed changes induced by specific cellular perturbations. To address this challenge, we established a semi-automated workflow to segment organellar membranes and reconstruct their underlying surface geometry in cryo-ET. To complement this workflow, we developed an open source suite of ultrastructural quantifications, integrated into a single pipeline called the surface morphometrics toolkit. This toolkit allows detailed mapping of spacing, curvature, and orientation onto reconstructed membrane meshes, highlighting subtle organellar features that are challenging to detect in three dimensions and allowing for statistical comparison across many organelles. To demonstrate the advantages of this approach, we combine cryo-ET with cryo-fluorescence microscopy to correlate bulk mitochondrial network morphology (i.e., elongated versus fragmented) with membrane ultrastructure of individual mitochondria in the presence and absence of endoplasmic reticulum (ER) stress. Using our toolkit, we demonstrate ER stress promotes adaptive remodeling of ultrastructural features of mitochondria including spacing between the inner and outer membranes, local curvature of the inner membrane, and spacing between mitochondrial cristae. We show that differences in membrane ultrastructure correlate to mitochondrial network morphologies, suggesting that these two remodeling events are coupled. Our toolkit offers opportunities for quantifying changes in organellar architecture on a single-cell level using cryo-ET, opening new opportunities to define changes in ultrastructural features induced by diverse types of cellular perturbations.

## INTRODUCTION

Cellular cryo-electron tomography (cryo-ET) is a powerful technique that enables high-resolution, three-dimensional (3D) views of protein and organellar structure in unfixed and fully hydrated conditions^1^. By combining cryo-focused ion beam (cryo-FIB) milling with automated data collection schemes^2, 3^, it has become routine to collect large datasets of thin, cellular three-dimensional reconstructions (tomograms) teeming with pristinely preserved proteins and organelles. Even as tremendous progress has been made in developing new algorithmic subtomogram averaging approaches to solve protein structures in cells^4–6^, it remains very challenging to quantitatively describe complex organellar membrane architectures (hereafter referred to as membrane ultrastructure) from cryo-tomograms. Organellar membrane remodeling plays a pivotal role in the cell’s ability to adapt and respond to various changes in the physiological state of the cell (e.g., of cellular stress), and is mediated through changes in the molecular interactions of proteins that reside within membrane compartments^7, 8^. As such, an improved ability to quantify organellar architectures from cryo-tomograms would both enable contextualization of distinct protein structures revealed by subtomogram averaging and reveal subtle, yet functionally relevant, changes in organellar ultrastructures associated with distinct cellular physiologies.

Despite the inherent 3D nature of cryo-tomograms, the two-dimensional (2D) virtual slices of cryo-tomograms are often used to quantify membrane ultrastructural parameters such as inter- and intra-membrane distances through manual designation of membrane boundaries, by locating distinct points along the membrane^9–12^. Although this can serve as a proxy to quantify 3D ultrastructure, this analysis is time-intensive and prone to both user-bias and inaccuracies in the precise location of membranes between 2D slices. Furthermore, the manual nature of this approach severely limits its applicability to perform ultrastructural quantification at a throughput and sample size sufficient to reveal substantial mechanistic insight. Towards a more accurate 3D representation, manual or automated^13^ voxel-based segmentation can be used to assign and label individual voxels within cryo-tomograms to specific cellular membranes^14^. Although qualitatively informative, voxel-based segmentation methods do not encode for parameters defining membrane geometry (i.e. connectivity), a property essential for robust quantifications of ultrastructural features like membrane curvature and spacing.

An alternative to using voxel segmentations to represent membranes is representing them as a meshwork of triangles. Triangle mesh surfaces represent the connectivity and implicit geometry of the membrane itself, enabling direct measurement of metrics like orientation and curvature. A recently developed approach enables more accurate and robust estimation of each triangle’s geometry, improving quantification of local curvature as well as the spatial relationship between individual triangles (i.e. membrane segments) both within the same (intra-) and between distinct (inter-) membrane compartments^15^. Despite these major advances, surface reconstructions using segmentations of membranes themselves (i.e. boundary segmentations) were not reliable, due to the incomplete (open) nature of the membrane segmentations limiting the effectiveness of the reconstruction algorithm^16^. In cryo-electron tomography, even perfect segmentations will never show complete and closed membranes, due to the substantial missing wedge^17^ making horizontal membranes invisible. To address these issues, quantifications using this new approach depended on using voxel segmentations that “fill” the entire internal volume encompassed within a membrane (i.e. compartment segmentations), which are not amenable to automation. Manual segmentation of filled membrane compartments is feasible for small and simple compartments, but the time intensive process severely limits throughput and, thus, the ability to aggregate ultrastructural quantifications across large tomographic datasets. The low throughput is further exacerbated in the context of complex, highly variable organellar membranes, which can be very challenging to accurately fill in. Thus, new strategies are required to improve throughput, automation, and quantification of membrane ultrastructures for application to cryo-electron tomography.

Mitochondria are an ideal target for the development of this approach. Mitochondria are highly pleomorphic organelles that function simultaneously as an interconnected network population and as discrete organellar units involved in energy production, ion homeostasis, cellular stress pathway integration, and innate immune signaling^18–20^. Their ability to perform these essential eukaryotic functions fundamentally depends on their dynamic remodeling both within the entire cellular mitochondrial network population (hereafter referred to as bulk mitochondrial morphology) and within their distinct outer and inner mitochondrial membranes (hereafter referred to as mitochondrial membrane ultrastructure)^21^. Changes to bulk mitochondrial morphology are mediated through the opposing processes of mitochondrial fusion and fission, and imbalances in these processes can promote interconnected (i.e. elongated) or disconnected (i.e. fragmented) networks that regulate pro-survival or pro-apoptotic mitochondrial functions^22, 23^. Within individual mitochondria, ultrastructures of both the outer and inner mitochondrial membranes (OMM and IMM, respectively) and the associations between these two membranes can exhibit dynamic remodeling to modulate metabolic and signaling functions^24^.

In particular, the IMM contains functional membrane folds, called cristae, whose ultrastructure is intimately linked to nearly all aspects of mitochondrial function. The architecture of the cristae is maintained through the activity and localization of several proteins (termed “shaping” proteins), including components involved in cellular respiration such as ATP synthase, whose dimerization at the matrix-associated cristae “tip” regions induces membrane curvature required for efficient mitochondrial respiration,^25–27^. Cristae shape is also maintained through specialized regions called cristae junctions, formed by the interaction of mitochondrial contact site and cristae organizing system (MICOS) and Optic Atrophy 1 (OPA1), which act as dynamic molecular staples to compartmentalize the distinct internal cristae environments and associate IMM cristae with the OMM^28–30^. Alterations to the expression, localization, or activity of these shaping proteins disrupt mitochondrial respiration and can promote the induction of apoptotic signaling through the release of cytochrome C^25, 31^. It is therefore important to understand the relationship between mitochondrial network morphology and membrane ultrastructure, and how they are altered in response to cellular perturbations.

Here, we sought to develop a correlative cryo-electron tomography workflow that enables quantitative analysis of complex membrane ultrastructures across distinct mitochondrial network morphologies and cellular physiologies with throughputs sufficient for statistical hypothesis testing. Specifically, we combined cryo-fluorescence microscopy with cellular cryo-ET imaging to correlate distinct mitochondrial network morphologies (e.g., elongated versus fragmented) to mitochondrial membrane ultrastructure. We developed and implemented a robust, semi-automated workflow^13^ to segment cellular membranes present in cryo-tomograms, and applied a screened Poisson reconstruction strategy^32^ to convert membrane voxel segmentations into implicit surface meshes that model complex cellular membrane geometries. To complement our surface generation workflow, we developed the surface morphometrics toolkit to measure several parameters of membrane shape including inter- and intra-membrane spacing, curvedness, and orientation across several hundred square microns of mitochondrial membranes. We applied this workflow to cells experiencing acute levels of endoplasmic reticulum (ER) stress, a condition known to promote adaptive remodeling of mitochondrial networks^33^, to explore how these changes influence membrane ultrastructure across distinct network morphologies. Our workflow for the first time allows us to distinguish statistically significant alterations in 3D ultrastructure in highly variable mitochondrial inner membranes.

## RESULTS

### Correlative cryo fluorescence microscopy and electron tomography enables association of bulk mitochondrial morphology with local membrane ultrastructure

We set out to understand the connection between the bulk morphology of the mitochondrial network within a single cell and the local membrane ultrastructure of the mitochondrial membranes within that same cell. We used cryo-fluorescence microscopy to image vitrified mouse embryonic fibroblasts whose mitochondria were labeled with a mitochondria-targeted GFP (MEF^mtGFP^), such that the morphology of each cell’s mitochondrial network could be readily assessed (Figure 1A-C, Figure 2A). We categorized cells for mitochondrial network morphology by blinded manual classification based on the fluorescence microscopy, then targeted cells with either fragmented or elongated mitochondrial networks for cryo-focused ion beam (cryo-FIB) milling to prepare thin (∼100-200 nm) lamellae (Figure 1D, Figure 2A). We collected tilt series of the mitochondria within these lamellae to directly correlate the cell’s bulk mitochondrial morphology (Figure 1C, Figure 2A) to the ultrastructure of mitochondrial membranes (Figure 1 F-H, Figure 2B). To further probe the connection between morphology and ultrastructure, we treated MEF^mtGFP^ cells with Thapsigargin (Tg), a small molecule that induces ER stress through inhibition of the sacro/ER Ca^2+^ ATPase (SERCA) and has been previously reported to promote mitochondrial elongation downstream of the ER stress-responsive kinase PERK through a process termed stress-induced mitochondrial hyperfusion (SIMH)^33^. We applied our correlative approach to identify and target specific Tg-treated MEF^mtGFP^ cells with either elongated or fragmented mitochondrial network morphologies for cryo-FIB milling and cryo-ET data acquisition and reconstruction. Visual inspection of the mitochondrial membranes within tomograms revealed variability in IMM architecture between mitochondria from different network morphologies and treatment groups, but some overall trends were apparent in elongated and fragmented mitochondrial populations in response to Tg-induced ER stress (Supplementary Figure 1). Notably, Tg treatment induced a shift towards regularly spaced disc-shaped (lamellar) cristae in the elongated population and towards a swollen tubular crista structure in the fragmented population. Due to the high degree of pleomorphism, with mixed populations in all conditions, we next established a workflow for ultrastructural quantification to determine the significance of these apparent changes.

**Figure 1.**
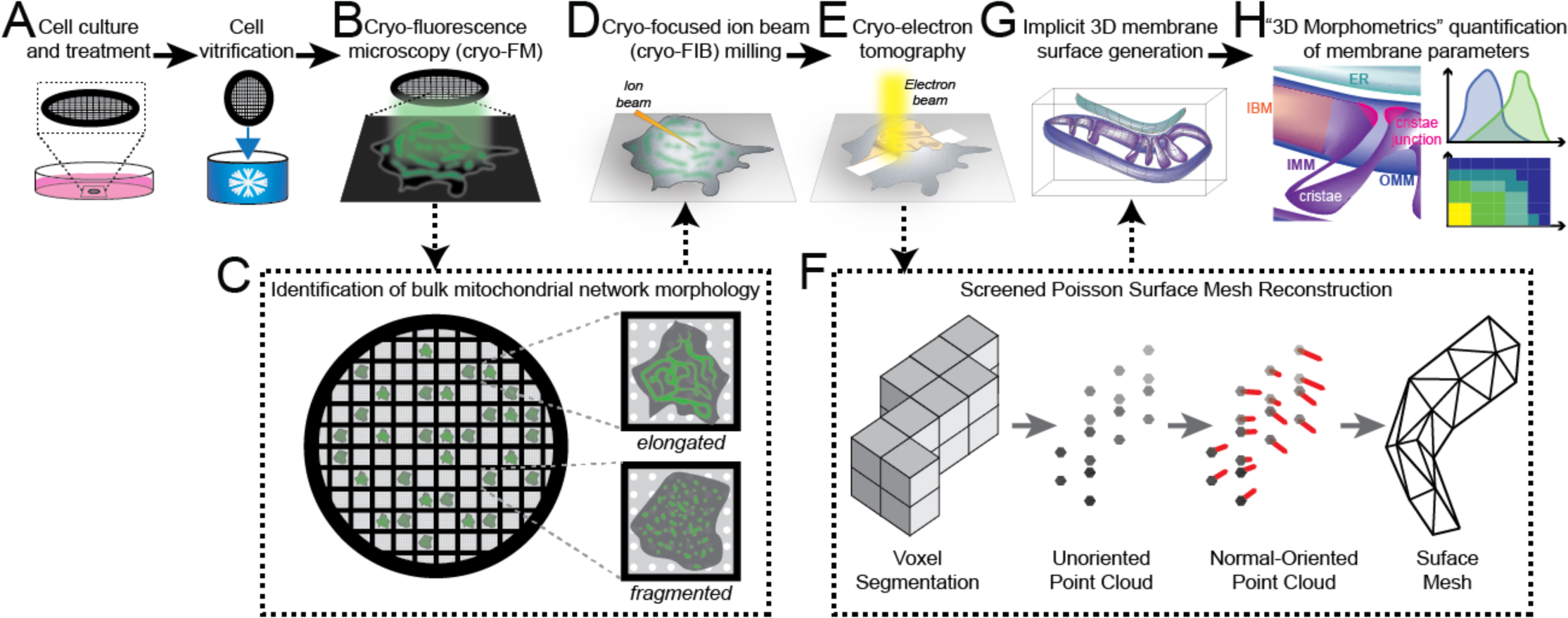
Correlative cellular cryo-electron tomography workflow for robust quantitative analysis of membrane ultrastructures between distinct mitochondrial network morphologies and treatment conditions. (A) MEF^mtGFP^ cells were cultured on transmission electron microscopy grids (black mesh circle) and treated with vehicle or thapsigargin (Tg, 500 nM) for 8 hours prior to vitrification via plunge freezing in ethane/propane mixture. (B) Vitrified MEF^mtGFP^ cells were imaged using cryo-fluorescence (cryo-FM). (C) Cryo-FM facilitated the identification of single cells with distinct elongated or fragmented bulk mitochondrial network morphologies. (D) These cells were targeted for cryo-focused ion beam (cryo-FIB) milling to generate thin sections (lamellae). (E) Lamella were imaged using standard cryo-ET imaging parameters to generate tilt series that were further reconstructed to generate three-dimensional (3D) cryo-tomograms. (F) Voxel segmentations labeling distinct cellular membranes were generated using a semi-automated approach^13^ prior to conversion to point clouds. Next, a normal-oriented vector is estimated for each point^61^. Normal-oriented point clouds were used to generate surface meshes using a novel application of a screened Poisson reconstruction method followed by masking to the original voxel segmentation^32, 61^. (G) Surface meshes model the implicit 3D ultrastructure of distinct organellar membranes. Surface coloring: inner mitochondrial membrane (IMM), purple; outer mitochondrial membrane (OMM), blue; endoplasmic reticulum (ER), teal. (H) Implicit geometries encoded within surface meshes are then used to perform 3D morphometrics to quantify several parameters that define membrane ultrastructure across mitochondria from distinct morphological and treatment conditions.

**Figure 2.**
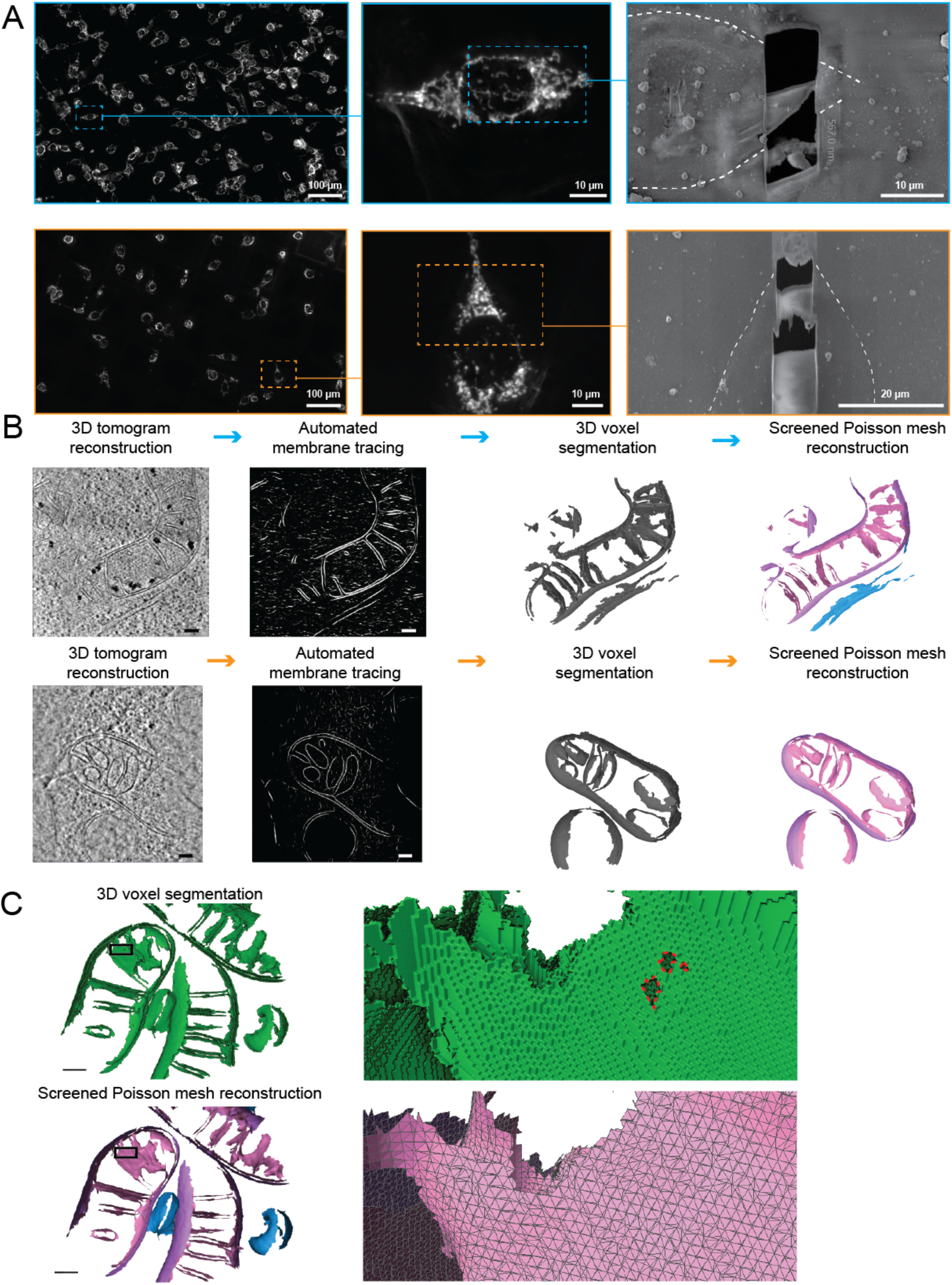
Application of correlative workflow to visualize cellular membrane ultrastructures in elongated and fragmented mitochondrial networks in MEF^mtGFP^ cells. (A) Identification by cryo-FM of elongated (top; cyan outline) and fragmented (bottom; orange outline) mitochondrial morphologies in MEF^mtGFP^ cells. Cryo-FM images of bulk mitochondrial morphology are then used for targeted cryo-FIB milling to generate thin lamellae of ∼150-200 nm. MEF^mtGFP^ cell periphery is outlined in dashed white line. (B) Virtual slides of tomograms of MEF^mtGFP^ cells containing elongated (top) and fragmented (bottom) mitochondria are traced using the automated tomosegmemtv program^13^, followed by manual cleanup using AMIRA software to generate 3D voxel segmentations. 3D voxel segmentations are then converted into implicit surface meshes using the screened Poisson mesh reconstruction. Scale bars = 100 nm. Surface coloring: IMM in pink, OMM in purple, ER in blue. (C) Detailed views of voxel (green) and triangle surface mesh (IMM: pink, OMM: purple, ER: blue) show the improved smoothness and hole-filling of surfaces generated with the screened Poisson mesh approach. Zoomed insets are rotated backwards 20° to highlight features. Holes are outlined in red on the voxel segmentation.

### Development of a framework to automate quantification of ultrastructural features of cellular membranes

In order to identify changes in different mitochondrial populations with and without induction of ER stress, we developed a semi-automated approach to quantify membrane features in three dimensions without the need for bias-prone manual measurements. We leveraged recent work^15^ that established a robust approach for estimating curvature in membranes and developed a framework for additional quantifications using triangular mesh models of membranes. However, the generation of high quality surface meshes has previously required labor-intensive manual segmentation approaches. These include manual contouring of membranes^34, 35^ and manual inpainting of entire compartments^15^, both of which are especially challenging with the complex and highly curved IMM and ER membrane. With these methods, it would be impractical to analyze a sufficient number of tomograms to overcome the variability between mitochondria observed in these distinct mitochondrial populations (elongated versus fragmented) and upon treatment (vehicle versus Tg). Instead, we developed a semi-automated workflow to enable analysis at higher throughput (Figure 1F, Figure 2B). We used a segmentation strategy that takes advantage of the tomosegmemtv program^13^, which identifies and enhances regions of the tomogram with membrane-like voxel values. This allowed manual classification of membrane identity into mitochondrial IMM and OMM and ER membrane, as well as manual cleanup of individual membrane segmentations using AMIRA software (Thermo Fisher Scientific). With this strategy, we were able to segment 32 tomograms containing mitochondria, divided between the elongated and fragmented bulk morphology populations and the two treatment groups (Figure 2, Supplementary Figure 1). In order to prepare high quality surface meshes appropriate for quantification, we established a fully automated surface reconstruction pipeline using the screened Poisson algorithm^32^ (Figure 1G, Figure 2B, Supplementary figure 1). This allowed reconstruction of much higher quality membrane surface as compared to previously developed approaches for mesh reconstruction from membrane segmentations (Figure 2C, Supplementary Figure 2, Supplementary Movie 1). These surfaces were then passed into the 3D surface morphometrics workflow, generating quantifications such as inter- and intra-membrane distances, membrane curvature, and relative membrane orientations (Figure 1H). This 3D surface morphometrics toolkit is available at https://github.com/grotjahnlab/surface_morphometrics.

### Quantification of IMM-OMM spacing in mitochondria identifies ultrastructural differences in elongated and fragmented mitochondrial networks

The most distinctive feature of mitochondria are the juxtaposed inner and outer mitochondrial membranes. Cryo-electron tomography is uniquely able to reveal the details of the architecture of these two membranes in 3D, and we sought out to use our tomograms to precisely measure their spacing. We measured the distance to the closest triangle on the IMM mesh for each triangle in the OMM mesh (Figure 3A). We then visualized the spatial distribution of these distances directly on the generated surface mesh reconstructions of the OMM of individual mitochondria visible within tomograms (Figure 3B). The resulting surface maps showed regions marked by the largest OMM-IMM distance representing the distribution of the crista junctions, but also revealed general increases in distance in the fragmented over elongated mitochondria in both vehicle- and Tg-treated conditions (Figure 3B). In order to determine how consistent these effects were across all of the segmented tomograms, we plotted a histogram of the combined distribution of distances for all surface mesh triangles within each condition (Figure 3C). This revealed an overall increase in inter-membrane distance in fragmented mitochondria. Furthermore, we observed a subtle decrease in the peak OMM-IMM distance after Tg-induced ER stress in elongated mitochondria, contrasted by a Tg-dependent increase in peak and variability of distances in fragmented populations. We then assessed the peaks of the distance distribution for each tomogram as independent observations and identified statistically significant increases in spacing in fragmented vs elongated cells, by 1.7 nm in the vehicle-treated condition and by 3.9 nm in the Tg-treated condition. Additionally, we confirmed that the 0.8 nm decrease in distance of the elongated mitochondria after Tg treatment was statistically significant, while the increase in spacing in fragmented cells after Tg treatment was not significant (Figure 3D), demonstrating the power of our approach to quantify and test the significance differences across distinct bulk mitochondrial network morphologies.

**Figure 3.**
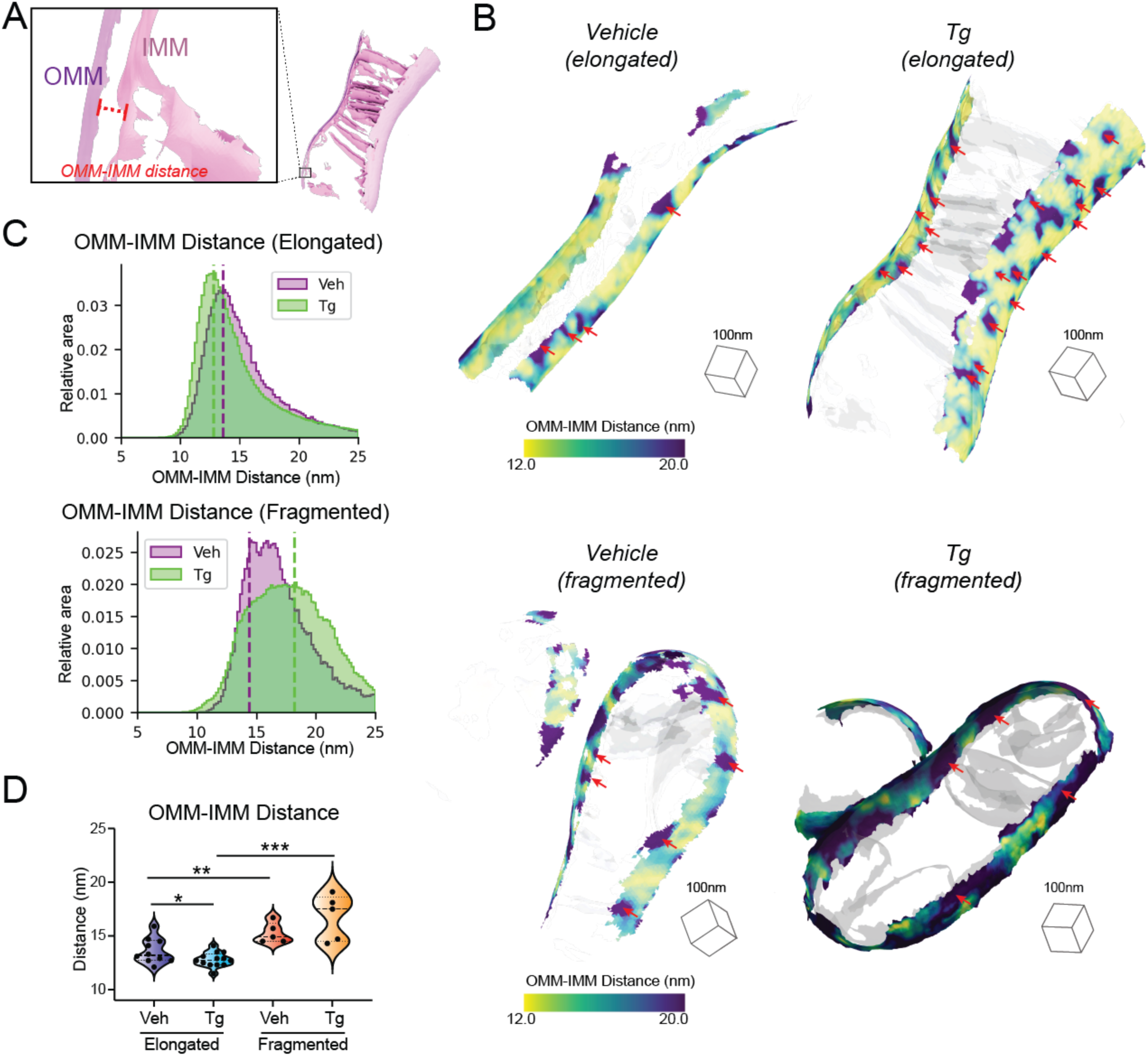
Inter-mitochondrial membrane distance is dependent on mitochondrial network morphology and presence or absence of ER stress. (A) Surface membrane reconstruction defining outer mitochondrial membrane (OMM, purple) and inner mitochondrial membrane (IMM, pink) distance measurement. (B) Representative membrane surface reconstructions of elongated (top) and fragmented (bottom) mitochondria in MEF^mtGFP^ cells treated with vehicle and thapsigargin (Tg; 500 nM, 8h). The OMM surface is colored by outer-to-inner (OMM-IMM) membrane distance, and the IMM surface is shown in transparent gray. Red arrows indicate regions on OMM with large OMM-IMM distances that correspond to cristae junctions. (C) Quantification of OMM-IMM distances of elongated (top) and fragmented (bottom) mitochondria in MEF^mtGFP^ cells treated with vehicle and Tg. (D) Quantification of peak histogram values from each tomogram within the indicated treatment and mitochondrial morphology class. Individual quantification from n=5-12 cryo-tomograms is shown. p-values from Mann-Whitney U test are indicated. *p<0.05; **p<0.01, ***p<0.005.

### Mitochondrial cristae and junction spacing in mitochondria are differentially impacted by Tg in elongated and fragmented mitochondrial networks

The IMM is divided between the inner boundary membrane (IBM), which is in close proximity to the OMM, and the invaginated mitochondrial cristae (Figure 4A), which are maintained as separate functional compartments by the organization of the MICOS complex and the protein OPA1 at crista junctions^29, 36^. We sought to understand how the organization of mitochondrial cristae and their junctions differed between fragmented and elongated mitochondria populations in the presence and absence of the ER stressor, Tg. We classified the IMM into three compartments (IBM, crista junctions, and crista bodies) based on their distance from the OMM such that quantifications could be performed based on the functional components of the membrane (Figure 4A). We then measured the distance between the membrane segments on either side of the crista bodies and junctions (intra-crista and intra-junction distances, respectively), as well as the distance between adjacent crista bodies and junctions (inter-crista and inter-junction distances, respectively) (Figure 4B). We quantified these effects as combined histograms showing the change in different populations upon treatment with Tg, and as violin plots of the histogram peaks for each tomogram (Figure 4C-F). Based on these distributions, we identified statistically significant but opposing changes in intra-crista body spacing between the elongated and fragmented morphologies induced by Tg treatment. In elongated mitochondria, crista spacing was reduced by 4.5 nm upon Tg treatment, while in fragmented mitochondria the spacing increased by 3.2 nm upon Tg treatment (Figure 4C). However, we did not observe significant changes in inter-cristae distance, intra-junction distance, or inter-junction distance (Figure 4D-F), with the exception of a 9.4 nm increase in intra-junction spacing in the Tg-treated condition of elongated mitochondria relative to fragmented mitochondria (Figure 4E), suggesting that ER stress promotes relaxation of the junction in fragmented mitochondria.

**Figure 4.**
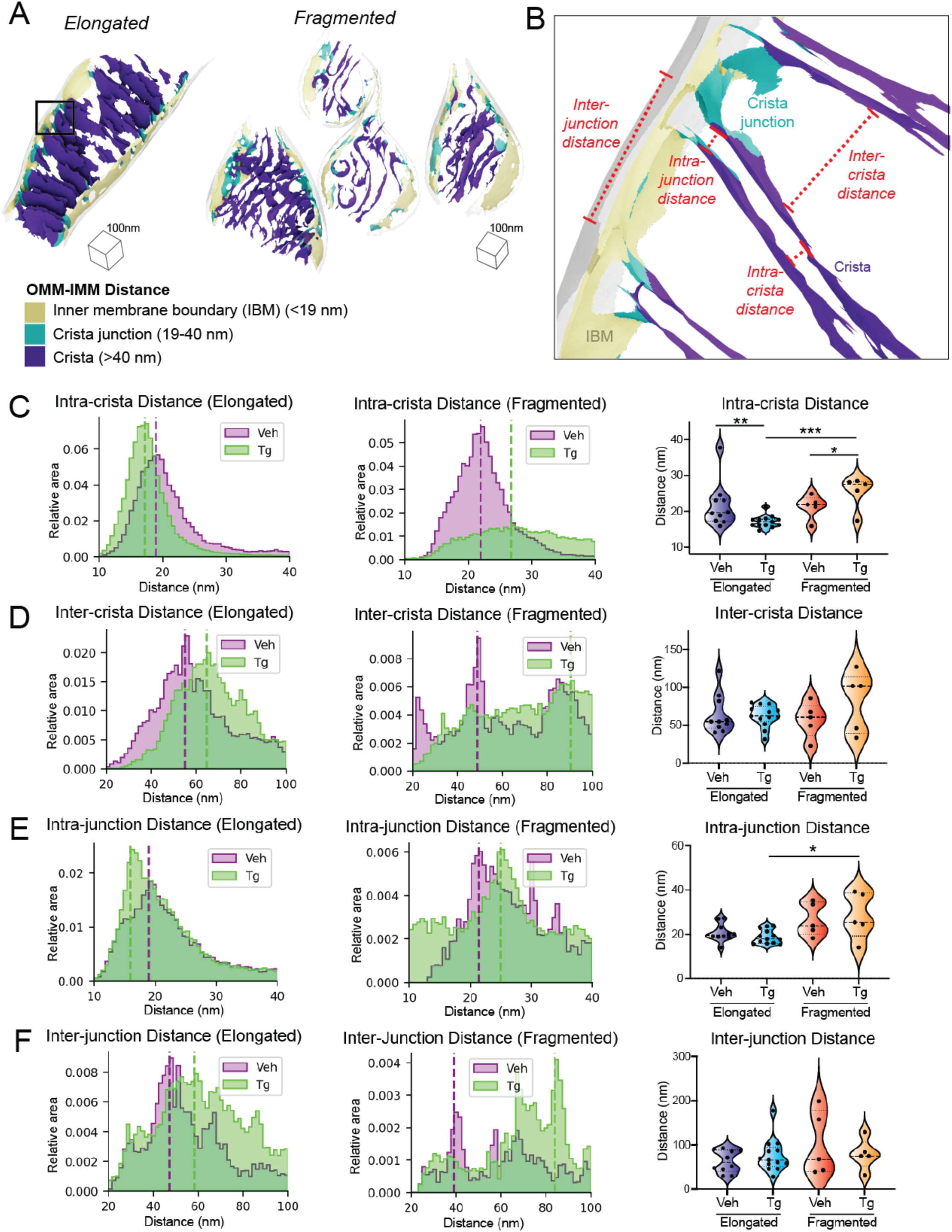
Spacing within and between mitochondrial cristae is differentially altered by Tg-induced ER stress in fragmented and elongated populations. (A) Representative membrane surface reconstructions of elongated (left) and fragmented (right) mitochondria in MEF^mtGFP^ cells treated with thapsigargin (Tg; 500 nM, 8h) showing subdivision of IMM compartments (inner membrane boundary (IBM), tan; junction, cyan; crista, purple) as defined by OMM-IMM distance. The OMM surface is represented as a transparent gray mesh. (B) Enlarged inset of boxed region in elongated example from (A) showing surface membrane reconstruction colored by inner membrane compartments defining intra- and inter-crista and junction measurements. (C) Quantification of intra-crista membrane distances of elongated (left) and fragmented (middle) mitochondria in MEF^mtGFP^ cells treated with vehicle and Tg. Graph (right) showing peak histogram values from each tomogram within each corresponding treatment and mitochondrial morphology class. Individual quantification from n=5-12 cryo-tomograms is shown. p-values from Mann-Whitney U test are indicated. *p<0.05; **p<0.01, ***p<0.005. (D) Quantification of inter-crista membrane distances of elongated (left) and fragmented (middle) mitochondria in MEF^mtGFP^ cells treated with vehicle and Tg. Graph (right) showing peak histogram values from each tomogram within each corresponding treatment and mitochondrial morphology class. Individual quantification from n=5-12 cryo-tomograms is shown. (E) Quantification of intra-junction membrane distances of elongated (left) and fragmented (middle) mitochondria in MEF^mtGFP^ cells treated with vehicle and Tg. Graph (right) showing peak histogram values from each tomogram within each corresponding treatment and mitochondrial morphology class. Individual quantification from n=5-12 cryo-tomograms is shown. p-values from Mann-Whitney U test are indicated. *p<0.05. (F) Quantification of inter-junction membrane distances of elongated (left) and fragmented (middle) mitochondria in MEF^mtGFP^ cells treated with vehicle and Tg. Graph (right) showing peak histogram values from each tomogram within each corresponding treatment and mitochondrial morphology class. Individual quantification from n=5-12 cryo-tomograms is shown.

### IMM curvedness is differentially sensitive to Tg treatment in elongated and fragmented mitochondrial networks

We next quantified the membrane curvedness locally at every mesh triangle position on each IMM surface to identify regions of local high and low curvature (Figure 5A)^15^. Visual inspection of these surfaces suggested notable decreases in curvedness in the flat and lamellar cristae observed in the Tg-treated elongated cells, while in cells with fragmented mitochondria, Tg appeared to increase curvedness in the swollen and tubular cristae (Figure 5A). To understand the degree to which these effects differed across populations and not just within individual mitochondria, we quantified the total curvedness of the IMM across different mitochondrial network morphologies and treatment conditions (Figure 5B). Surprisingly, despite apparent visual difference, no significant changes were observed in IMM curvedness overall (Figure 5B). Because visual inspection revealed local changes in curvature in the cristae and junctions, rather than uniformly across the IMM, we broke down these quantifications into the IMM’s functional subcompartments (Figure 4A, Figure 5C-E). Within the IBM, we observed a significant decrease in curvedness in fragmented relative to elongated mitochondrial populations in cells treated with vehicle (Figure 5C). Further, we found that Tg treatment significantly increased curvedness in elongated mitochondria, while no significant increase was observed in fragmented mitochondria from Tg-treated cells. This led to a significant increase in IBM curvedness in elongated mitochondria over fragmented mitochondria in Tg-treated cells (Figure 5C). Since the IBM’s curvedness is constrained to be near-identical to that of the OMM, this metric likely reports primarily on the radius of the mitochondria as a whole, suggesting that fragmented mitochondria are larger in radius despite their apparent shorter overall length, while ER stress slightly reduces the radius of elongated mitochondria (Figure 5A, Supplemental Figure 1).

**Figure 5.**
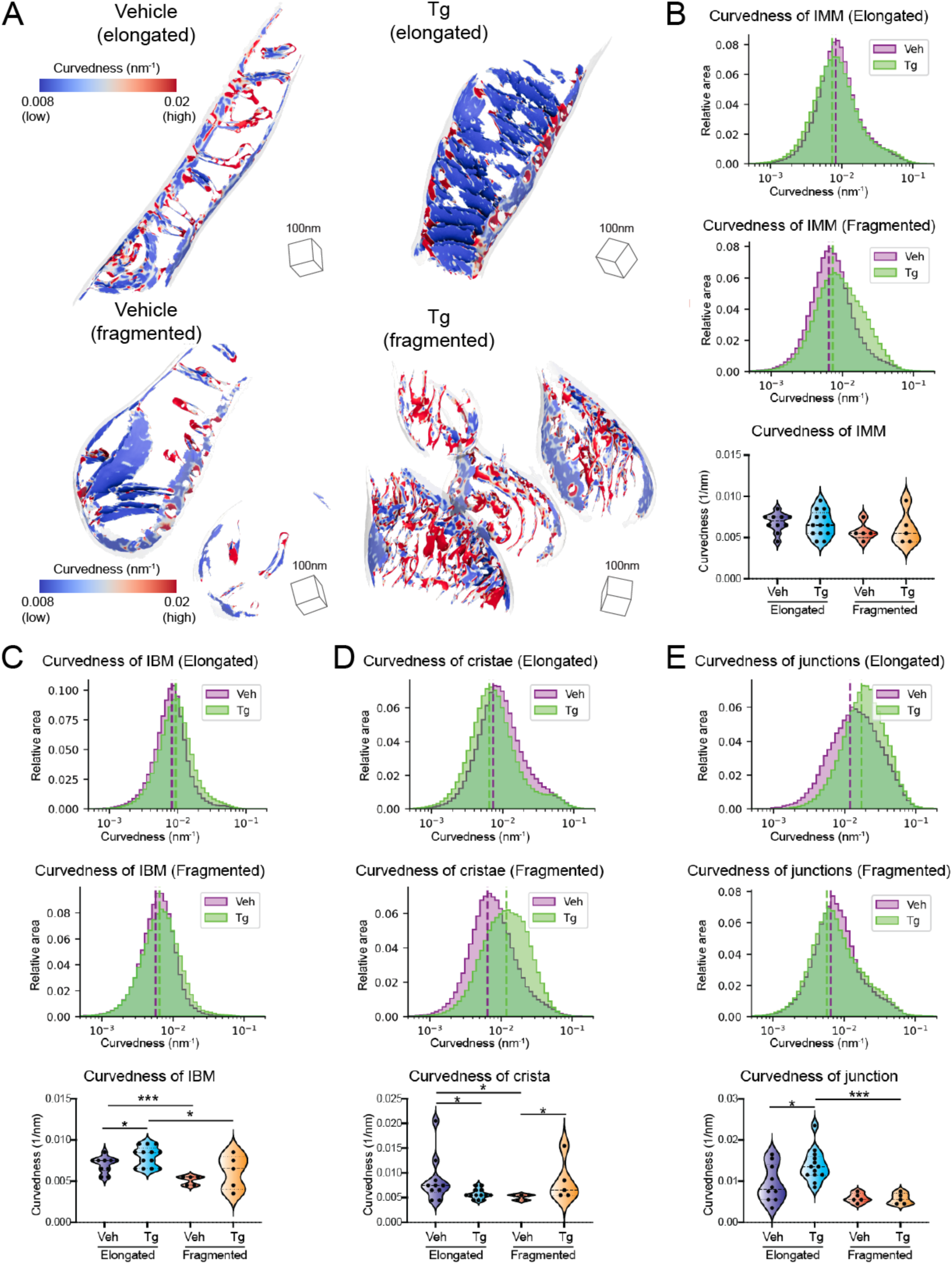
The curvedness of mitochondrial cristae and crista junctions are dependent both on mitochondrial network morphology and Tg-induced ER stress. (A) Representative membrane surface reconstructions of elongated (top) and fragmented (bottom) mitochondria in MEF^mtGFP^ cells treated with vehicle and thapsigargin (Tg; 500 nM, 8h). The IMM surface is colored by membrane curvedness and the OMM surface is represented as a transparent gray mesh. Curvedness is an unsigned combination of the two principle components of curvature and is used because the surfaces normal vectors do not have canonical sign. (B) Quantification of total IMM curvedness of elongated (top) and fragmented (middle) mitochondria in MEF^mtGFP^ cells treated with vehicle and Tg. Graph (bottom) showing peak histogram values from each tomogram within each corresponding treatment and mitochondrial morphology class. Individual quantification from n=5-12 cryo-tomograms is shown. (C) Quantification of inner boundary membrane (IBM) curvedness of elongated (top) and fragmented (middle) mitochondria in MEF^mtGFP^ cells treated with vehicle and Tg. Graph (bottom) showing peak histogram values from each tomogram within each corresponding treatment and mitochondrial morphology class. Individual quantification from n=5-12 cryo-tomograms is shown. p-values from Mann-Whitney U test are indicated. *p<0.05, ***p<0.005. (D) Quantification of cristae curvedness of elongated (top) and fragmented (middle) mitochondria in MEF^mtGFP^ cells treated with vehicle and Tg. Graph (bottom) showing peak histogram values from each tomogram within each corresponding treatment and mitochondrial morphology class. Individual quantification from n=5-12 cryo-tomograms is shown. p-values from Mann-Whitney U test are indicated. *p<0.05. (E) Quantification of cristae junction curvedness of elongated (top) and fragmented (middle) mitochondria in MEF^mtGFP^ cells treated with vehicle and Tg. Graph (bottom) showing peak histogram values from each tomogram within each corresponding treatment and mitochondrial morphology class. Individual quantification from n=5-12 cryo-tomograms is shown. p-values from Mann-Whitney U test are indicated. *p<0.05, ***p<0.005.

In contrast, larger changes were observed in cristae and crista junctions. The cristae of fragmented mitochondria were less curved than those of elongated mitochondria in the absence of ER stress (Figure 5D). Interestingly, Tg treatment induced significant but opposing changes in fragmented and elongated mitochondria. In elongated mitochondria, curvedness of cristae decreased upon Tg treatment, while in fragmented mitochondria, curvedness increased upon Tg treatment. In crista junctions, the most notable difference was an increase in curvedness in Tg-treated elongated cells compared to both the vehicle-treated elongated mitochondria and the Tg-treated fragmented mitochondria (Figure 5E). This reflects and confirms the visual observation of crista ordering upon Tg treatment in elongated mitochondria in the tomograms.

### Membrane orientations in elongated and fragmented mitochondria differ upon Tg treatment

We next sought to understand to what degree the orientation of mitochondrial cristae were regulated, and how they changed in different mitochondrial morphologies as well as in response to Tg-induced ER stress. We took advantage of the 3D nature of our surface reconstructions to measure the orientation of each triangle in the surface mesh relative to the nearest point on the OMM, as well as to the cell’s growth plane (Figure 6A). Visual inspection revealed that the highly organized lamellar phenotype that was typical of elongated mitochondria in Tg treated cells was almost entirely held orthogonal to the OMM, forming the distinctive ladder-like phenotype (Figure 6B). Similarly, most but not all of these lamellar cristae were remarkably vertical relative to the growth plane of the cell (Figure 6C). We plotted the angles of the cristae and junctions relative to the OMM and the growth plane, revealing changes primarily in the variance of the angles rather than the peak position. Because of this, we quantified the standard deviation of these values for each tomogram in order to assess statistical significance of observed changes (Figure 6D-G). This revealed a significant decrease in the variability of the angle of the crista bodies relative to the OMM for elongated mitochondria in Tg-treated cells (Figure 6D). We also identified a significant decrease in variability of the verticality of the crista body in Tg-treated elongated relative to fragmented mitochondria (Figure 6F). In both cases, these differences corresponded to increased control over the orientation of the ladder-like lamellar cristae observed in the elongated Tg-treated condition. Despite the changes in crista orientation, no significant changes were observed in the angle of junctions relative to the OMM (Figure 6E) or the growth plane (Figure 6G).

**Figure 6.**
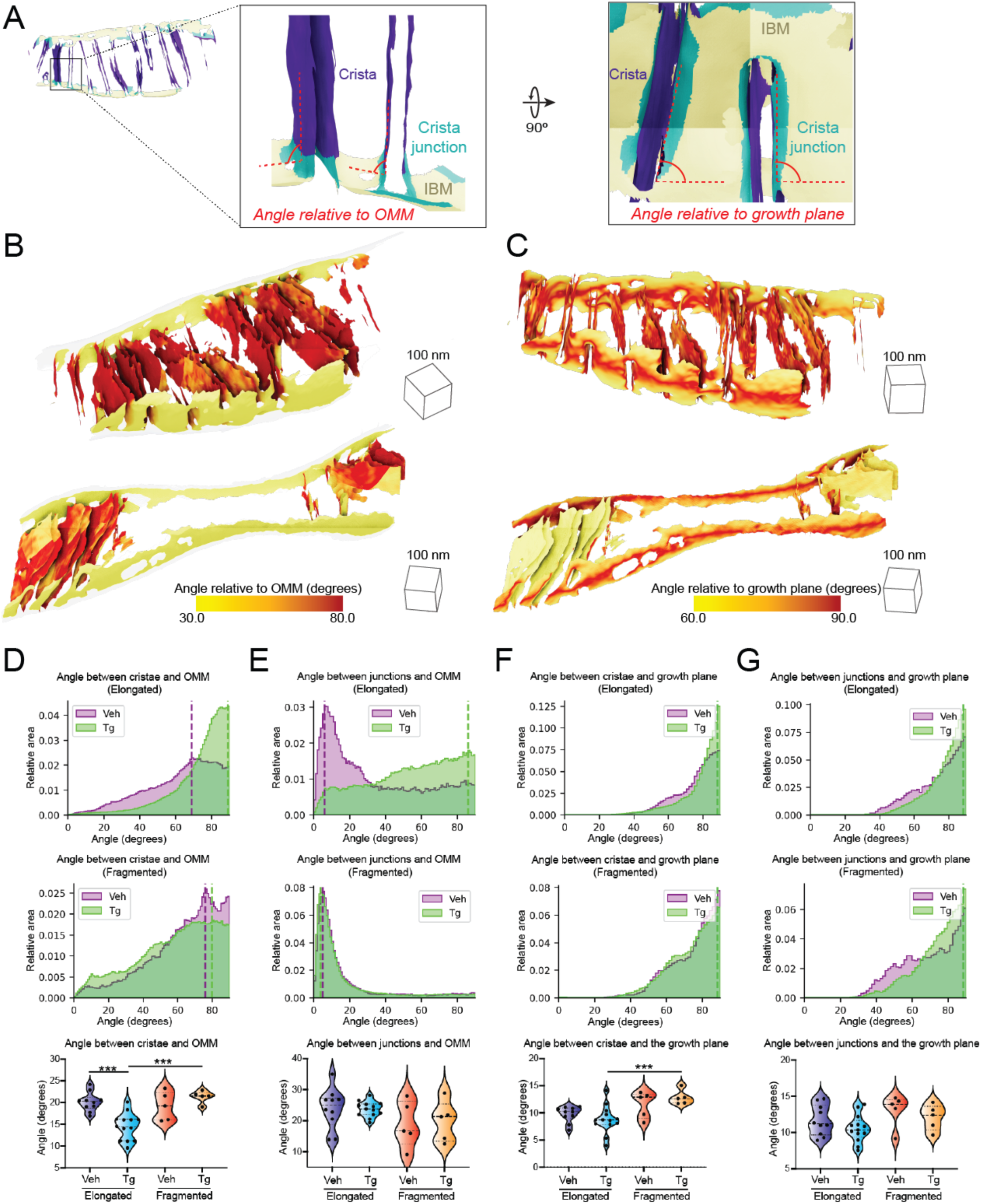
Tg-induced ER stress drives changes in crista orientation in elongated mitochondrial networks. (A) Surface membrane reconstruction colored by inner membrane compartments (inner membrane boundary (IBM), tan; junction, cyan; crista, purple;) defining angular measurements, between the surface and the nearest OMM or to the growth plane of the cell. (B) Representative membrane surface reconstruction of a rigid and lamellar Tg-treated elongated mitochondrion, colored by angle of IMM relative to OMM. (C) Representative membrane surface reconstruction of a less rigidly oriented Tg-treated elongated mitochondrion, colored by angle of IMM relative to the growth plane of the cell. (D) Quantification of angle of cristae relative to OMM in elongated (top) and fragmented (middle) mitochondria in MEF^mtGFP^ cells treated with vehicle and Tg. Graph (bottom) showing distribution of standard deviations within each corresponding treatment and mitochondrial morphology class. Individual quantification from n=5-12 cryo-tomograms is shown. p-values from Mann-Whitney U test are indicated. ***p<0.005. (E) Quantification of angle of junctions relative to OMM in elongated (top) and fragmented (middle) mitochondria in MEF^mtGFP^ cells treated with vehicle and Tg. Violin plot (bottom) showing distribution of standard deviations within each corresponding treatment and mitochondrial morphology class. Individual quantification from n=5-12 cryo-tomograms is shown. (F) Quantification of angle of cristae relative to growth plane in elongated (top) and fragmented (middle) mitochondria in MEF^mtGFP^ cells treated with vehicle and Tg. Graph (bottom) showing distribution of standard deviations within each corresponding treatment and mitochondrial morphology class. Individual quantification from n=5-12 cryo-tomograms is shown. p-values from Mann-Whitney U test are indicated. ***p<0.005. (G) Quantification of angle of junctions relative to growth plane in elongated (top) and fragmented (middle) mitochondria in MEF^mtGFP^ cells treated with vehicle and Tg. Graph (bottom) showing distribution of standard deviations within each corresponding treatment and mitochondrial morphology class. Individual quantification from n=5-12 cryo-tomograms is shown.

### Elongated and fragmented mitochondrial networks show differences in ER-mitochondrial contacts

Mitochondria-ER contact sites are sites of significant exchange of lipids and proteins have been previously associated with a wide variety of critical regulatory functions for mitochondria, ranging from mitochondrial fission^37^ to autophagy^38^. To investigate whether the remodeling we observed in response to Tg-induced ER stress was driven by alterations to the ultrastructure of these contact sites, we leveraged our 3D morphometrics workflow to quantify changes in ER-mitochondrial contacts in elongated vs fragmented mitochondrial networks as well as in ER stress induced by Tg. Distances between different organelles can be as readily calculated as distances within mitochondria, with the only additional challenge that not all tomograms contained ER, and not all ER present was interacting with mitochondria. We measured the distance between the OMM and the nearest ER in all tomograms where the ER was present, and visualized the contact sites on the surfaces of the OMM (Figure 7A-B). This showed detailed mapping of these functionally critical interactions in three dimensions. We also quantified the distribution of ER-OMM distances as a combined histogram (Figure 7C). None of the conditions were statistically significant, suggesting that changes in ER-mitochondrial contact distance or area at the imaged time-point were not driving the altered mitochondrial ultrastructure (Figure 7D).

**Figure 7.**
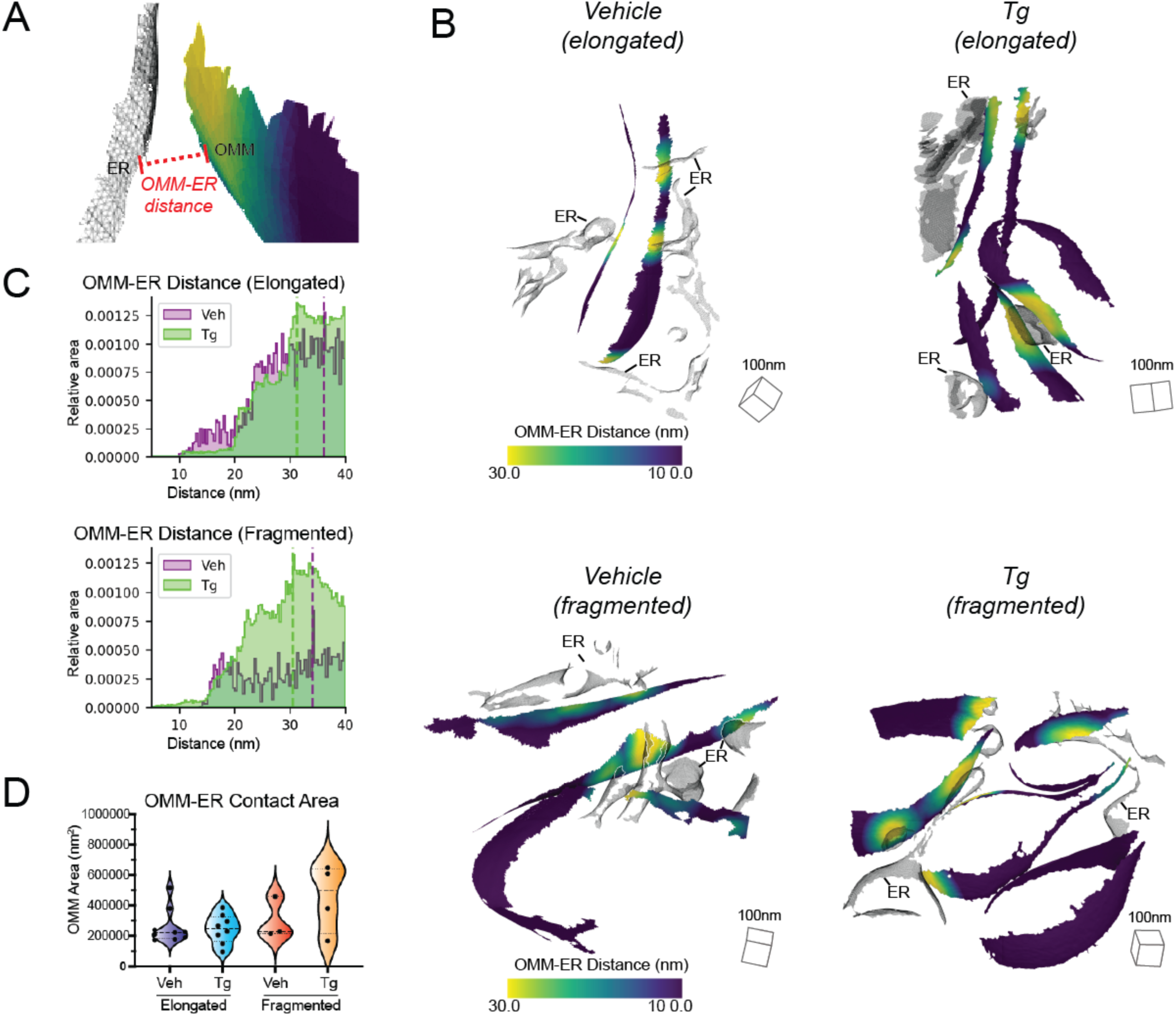
Mitochondria-ER contact sites can be localized in three dimensions on the outer membrane. (A) Surface membrane reconstruction defining OMM (purple) and ER (transparent) distance measurement. (B) Representative membrane surface reconstructions of elongated (top) and fragmented (bottom) mitochondria in MEF^mtGFP^ cells treated with vehicle and Thapsigargin (Tg; 500 nM) colored by OMM to ER (OMM-ER) membrane distance. (C) Quantification of OMM-ER distances of elongated (top) and fragmented (bottom) mitochondria in MEF^mtGFP^ cells treated with vehicle and Tg. (D) Graph showing peak histogram values from each tomogram within each corresponding treatment and mitochondrial morphology class. Individual quantification from n=5-9 cryo-tomograms is shown.

## DISCUSSION

We describe a correlative cellular cryo-electron tomography workflow for quantifying differences in membrane ultrastructure between distinct mitochondrial network morphologies and cellular physiology conditions. Our approach takes advantage of the distinct capabilities of multiple imaging modalities (fluorescence microscopy and cryo-ET) to correlate single-cell, organellar network morphologies to single-organellar membrane ultrastructures in varying cellular conditions. To our knowledge, this is the first application of a correlative approach that directly assesses how distinct organellar morphologies (e.g., elongated versus fragmented mitochondria) and cellular stress states (e.g., presence or absence of Tg-induced ER stress) influences mitochondrial membrane remodeling at nanometer-scale resolution. Our workflow can be divided into two functional steps: (1) single-cell targeting and (2) 3D surface morphometrics analysis (Figure 1, Figure 2). The first step involves identifying cells that exhibit a particular mitochondrial network morphology by cryo-fluorescence microscopy, targeting these cells for sample thinning to generate cellular lamellae by cryo-FIB milling, and imaging these cellular lamellae by cryo-electron tomography to reveal organellar membrane ultrastructures (Figure 1A-D, Figure 2A). The second step involves adapting semi-automated segmentation and surface mesh reconstruction algorithms to cryo-ET data, thereby enabling unbiased quantification of several membrane parameters at high-precision (Figure 1E-H, Figure 2B,C, Supplementary Movie 1). We determined that the screened Poisson reconstruction approach^32^ provides a more accurate estimation of the implicit membrane geometry of complex membranes, and can be used to model and quantify complex membranes such as the IMM (Supplementary Figure 2, Supplementary Movie 1).

To demonstrate proof-of-concept of our correlative workflow, we asked whether bulk changes in mitochondrial network morphologies were associated with ultrastructural remodeling of mitochondrial membranes, both in the presence and absence of ER stress. We identified cells with distinct mitochondrial morphologies (e.g., fragmented versus elongated) in both treatment conditions, and quantified mitochondrial membrane parameters such as inter- and intra-mitochondrial membrane distances (Figures 3-4), curvature (Figure 5), and orientation (Figure 6). We demonstrate that our approach can be used to identify changes to membrane ultrastructure in two ways. First, we can evaluate ultrastructural changes on an individual mitochondrion basis by overlaying values directly on individual triangles within our membrane surface maps (Figures 3B, 4A,B, 5A, 6B,C). This enables contextualization of membrane quantifications based on the specific membrane microenvironment. Second, we combine ultrastructural quantifications across many mitochondria within specific morphological or cellular physiological groups, thereby enabling us to establish statistical criteria for identifying trends even in the context of mitochondrial pleomorphism across all conditions (Figures 3C,D, 4C-F, 5C-E, 6D-G).

We first used our workflow to quantify the inter-membrane distance between the inner and outer membranes of mitochondria (Figure 3). These two membranes are mechanically linked by interactions between proteins on the two membranes^36, 39, 40^, and in some instances, there are reports of inner membrane-bound proteins acting in trans on the outer membrane^41^. The uniquely high resolution and 3D information afforded by cryo-ET allowed us to visualize the fluctuations of distances between these membranes (Figure 3B) as well as to quantify changes in the overall distributions (Figure 3C-D). We identified significant increases in the IMM-OMM spacing of mitochondria in fragmented networks over those in elongated networks. This increase in spacing may reduce the effectiveness of protein interactions across the two membranes, thereby increasing isolation of the IMM from external signals. In contrast, within the elongated population, Tg treatment reduced IMM-OMM spacing, suggesting tighter interactions that may be critical for the protective function of stress-induced mitochondrial hyperfusion.^33^ The difference in distance, from 13.6 to 12.8 nm between the centers of the membranes, is small, but correcting for the approximately 7 nm thickness of mitochondrial membranes^42^ yields a change in intermembrane contact distance from 6.6 to 5.8 nm, or a 15% reduction. This could have a significant effect on the potential for transmembrane activity of proteins, such as PISD, an IMM protein previously suggested to act on the OMM.^41^ The ability to consistently and robustly resolve such small but physiologically significant differences in intermembrane spacing opens new avenues for understanding how different perturbation may impact inter-membrane communication in mitochondria. The spatially resolved high OMM-IMM distance proximal to crista junctions may also eventually prove to be an effective automated localization signal for particle picking and guided subtomogram averaging of proteins, such as the TOM complex^36^, which associate with the MICOS complex and may impact MICOS complex organization^43^.

Mitochondrial inner membranes are functionally divided between the IBM and the mitochondrial cristae, with the cristae junctions that divide the two components organized by the MICOS complex^43^. We were able to automatically classify our inner membrane surface into the IBM, cristae, and junctions based on distance from the OMM, and using this classification were able to quantify specific architectural changes in cristae and junctions in different mitochondrial populations in the presence or absence of Tg-induced ER stress. We measured the membrane spacing within and between cristae and junctions (Figure 4), quantified the curvedness of the membranes (Figure 5), and measured their orientation relative to the OMM and the growth plane (Figure 6). We observed consistent but opposing changes between fragmented and elongated mitochondrial populations when exposed to Tg. In elongated populations of cells treated with Tg, cristae became narrower, less curved, and more vertical lamellar structures that were highly orthogonal to the OMM. This is consistent with previous reports of ultrastructural changes induced in brown adipocytes subjected to cold stress.^44^ In these conditions, modulations to the MICOS subunit Mic 19 remodels cristae architecture through the activity of the ER-stress responsive PERK kinase - a mechanism similar to that responsible for ER-stress induced mitochondrial elongation^33^. While future work is needed to determine whether cristae remodeling and mitochondrial elongation are regulated by the same mechanism, our results highlight that these two changes in mitochondrial shape are correlated in response to Tg-induced ER stress.

In contrast, cristae in fragmented populations became more swollen upon induction of ER stress, as measured by increases in intra-crista membrane spacing and crista curvature. Of particular interest, our workflow will enable local analysis of individual cristae on the basis of their spacing, curvature, and orientation to isolate different functional microenvironments that can be studied for differences in protein composition and conformation. Crista ultrastructure is largely controlled by MICOS^28^, ATP synthase^27^, and Opa1^29^, and we predict that the changes observed are driven by changes in composition, localization, or conformation of these proteins. Cryo-ET offers the unique combination of cellular and structural detail to pursue studies of how these proteins differ in distribution and conformation in different cristae even within the same mitochondrion, and how that is reflected in differences in mitochondrial architecture and function. Our workflow will facilitate future studies on the critical importance of mitochondrial remodeling induced by different types of cellular perturbations.

A notable caveat of the methodology we have developed and presented is the potential for bias in which components of membrane are segmented and which are not. In addition to the components of membrane that are severed during the cryo-FIB-milling process, membranes fade and disappear when their normal vectors fall into the missing wedge of angles not collected in the tilt series^1^, which can be seen in the lack of orientations less than 30° relative to the growth plane (Figure 6F,G). Additionally, the segmentation algorithm^13^ used in this study traces lower curvature membranes more easily than high curvature membranes, leading to limited segmentations of tight, high curvature crista junctions and crista tips. We have mitigated these biases to the best of our ability by comparing data collected in similar ways and processed identically across different populations, and we anticipate that improvements to both data collection and data processing will further reduce these biases in the future. The majority of these data were collected without the use of a post-column energy filter^45^ or phase plate^46^, both of which improve contrast of visible features within tomograms. Additionally, there have been recent advances in algorithmic approaches to improve the quality and completeness of segmentations^14, 47–49^. The flexibility of the screened Poisson surface reconstruction workflow will enable easy adaptation of these morphometric approaches to new segmentation tools.

Beyond incorporating new advances for ultrastructure, the automated surface reconstruction algorithm established in this paper for quantitative ultrastructural analysis presents additional potential for use in visual proteomics^50^ and subtomogram averaging. Finding proteins in noisy tomograms is challenging, and membrane segmentations have proven effective guides for localizing protein density that would otherwise be difficult to detect with traditional template matching approaches^51, 52^. These approaches have in the past depended on voxel segmentations, but moving to high quality meshes has the potential to improve particle picking further by constraining the orientation search to match the geometry of the membrane, as well as providing a more consistent central position than voxel segmentations which might have variable thickness. Prior knowledge of orientation has proven to be valuable for achieving successful initial alignments for subtomogram averaging as well^53^. Membrane meshes have been used very successfully to guide subtomogram averaging in the past^35, 54^, but these have relied on manual segmentation and thus have been low throughput and restricted to relatively simple geometry. Surfaces generated automatically with our software have the potential to extend this guided approach to larger numbers of tomograms and to much more complex biological environments.

There is tremendous potential for cryo-ET to enable better understanding of function and regulation by associating changes in protein conformation with the local cellular milieu^26, 51, 55–60^. Cryo-ET’s ability to reveal both 3D protein conformation and the cellular ultrastructures surrounding it allows connections between the two that cannot be determined any other way. By using correlative light microscopy approaches to incorporate knowledge of the physiological state of the cell in combination with detailed local membrane morphometrics, it will be possible to gain a much more detailed understanding of how both global cellular state and local membrane ultrastructure correlate to protein localization and conformation. Moving from qualitative association to quantification with the measurement of statistical significance, as outlined here, will improve the power of structure-context mapping to understand how protein structure and function impact complex cellular processes.

## ACKNOWLEDGEMENTS

We thank Bill Anderson and Mengyu Wu at The Scripps Research Institute electron microscopy facility for microscope support, and Jean-Christophe Ducom at The Scripps Research Institute for computational support.

We thank Maria Salfer and Antonio Martinez-Sánchez for software support. We also thank David DeRosier, Laura Newman, Hamid Rahmani, and Linda Joosen for their critical input on the manuscript. D.A.G. is supported by the Nadia’s Gift Foundation Innovator Award of the Damon Runyon Cancer Foundation (DRR-65-21). B.A.B. is supported by an American Cancer Society postdoctoral fellowship (PF-21-075-01-CCB). R.L.W. is supported by the National Institutes of Health (NIH) grant R01NS095892.

## MATERIALS AND METHODS

### Preparation of vitrified mouse embryonic fibroblasts on cryo-EM Grids

Mouse embryonic fibroblasts with mitochondria labeled GFP (MEF^mtGFP^)^62^ were cultured in Dulbecco’s Modified Eagle Medium + GlutaMAX (Gibco) additionally supplemented with HiFBS (10%) and glutamine (4 mM) on fibronectin-treated (500 ug/ml, Corning) and UV sterilized R ¼ Carbon 200-mesh gold electron microscopy (EM) grids (Quantifoil Micro Tools). After 15-18 hours of culture, MEF^mtGFP^ cells were transferred to fresh media supplemented with DMSO in the vehicle treatment group, or fresh media supplemented with Thapsigargin (500nM, Fisher Scientific) in the Tg treatment group. After 8 hours of incubation, samples were plunge-frozen in a liquid ethane/propane mixture using a Vitrobot Mark 4 (Thermo Fisher Scientific) with manual back-blotting.

### Cryo-fluorescence microscopy and Mitochondria Network Morphology Scoring

Fluorescence and bright-field tiled image maps (atlases) of EM grids containing vitrified cellular samples were acquired with a Leica CryoCLEM microscope (Leica) were collected using Leica LAS X software (25 um Z stacks with system optimized steps, GFP channel ex: 470, em : 525). Z stacks were stitched together in LAS X navigator to provide a single mosaic of maximum projections of the GFP signal, enabling rapid identification of the bulk mitochondrial morphology for each cell. For classification of mitochondrial network morphologies, max projections of individual tiles within fluorescence atlases of MEF^mtGFP^ cells were randomized and blinded. Five researchers classified the cells as containing primarily elongated or fragmented mitochondria (see Figure 2a for examples of scoring). Atlases were then exported from LAS X and loaded into MAPS (3.13) (Thermo Fisher) for fluorescence guided milling.

### Fluorescence Guided Milling

Cryo-Focused Ion Beam milling of lamellae was performed using an Aquilos dual-beam cryo-FIB/SEM instrument (Thermo Fisher Scientific). The fluorescence atlases were overlaid and aligned to an SEM atlases of the same grid to target milling of MEF^mtGFP^ cells with distinct mitochondrial network morphologies, as determined during blind classification (described above). MEF^mtGFP^ cell targets were chosen based on their position within grid squares, the thickness of the ice in their vicinity, and based on their bulk morphology as assessed by the GFP fluorescence channel. Prior to milling, EM grids were first coated with a thin platinum sputter, then coated with an organometallic platinum layer using a gas injection system (GIS) for 3-4 seconds using an automation script^63^, followed by a third and final layer of platinum sputter was added. Targeted cells were milled manually^64^ or using an automated cryo preparation workflow^2^ both methods used xT software with MAPS (Thermo Fisher). Minimal SEM imaging for monitoring was done at 2keV to ensure thin lamella generation while avoiding radiation damage. A total of 6 treated grids and 8 untreated grids were milled for further tomography analysis.

### Tilt Series Data Collection

EM grids containing lamellae were transferred into a 300keV Titan Krios microscope (Thermo Fisher Scientific), equipped with a K2 Summit direct electron detector camera (Gatan). A small subset of data was collected after the installation of a BioQuantum energy filter (Gatan). Individual lamellae were montaged with low dose (1e^-^/Å^2^) at high magnification (53,000x nominal magnification) to localize cellular regions containing mitochondria, which were identified by their distinctive inner and outer mitochondrial membranes. Data was collected to maximize the number of non-overlapping fields of view containing mitochondria, with no targeting of specific observed membrane ultrastructure. Tilt series were acquired using SerialEM software (Mastronarde, 2005) with 2 steps between -60 and +60. Data was collected with pixel sizes of 3.11Å, 3.35 Å, or 3.55 Å, and a nominal defocus range between -5 to -10 µm. A subset of data was collected with dose fractionation, with 10 0.1e/Å^2^ frames collected per second. The total dose per tilt was 0.9-1.2 e/Å^2^, and the total accumulated dose for the tilt series was under 70 e/Å^2^.

### Tilt series processing and reconstruction

Dose fractionated tilt series micrograph movies underwent CTF estimation and motion correction in Warp and combined into averaged tilt series for alignment. Both non-fractionated and fractionated tilt series then were aligned using patch tracking in etomo^65^, with 4 times binning and 400 binned pixel patches. Resulting contours were manually curated to remove poorly aligning patches, and the remaining contours were used for alignment and reconstruction with weighted back projection into 4 times binned tomograms. Tomogram thicknesses ranged from 95 nm to 233 nm.

### Semi-automated Segmentation

Automated detection and enhancement of mitochondrial membranes was performed using the TomoSegMemTV software package^13^. Settings were optimized for individual tomograms to maximize the membrane quality. Typical parameters for membrane enhancement were as follows:

- scale_space -s 3
- dtvoting -s 12
- surfaceness -m 0.17 -s 0.8 -p 0.8
- dtvoting -w -s 10
- surfaceness -S -s 0.75 -p 0.75 -l 20
- thresholding -l 0.01 -2 30
- global analysis -3 100

The output correlation volumes containing high pixel intensity corresponding to membrane locations were imported into AMIRA (Thermo Fisher) for manual curation. Mitochondrial inner and outer membranes and endoplasmic reticulum were segmented and designated as separate labels in Amira using 3D threshold-based selection and the 3D magic wand tool, and very small gaps in segmentation were filled in manually using a paint tool.

### Automated surface reconstruction

The process for conversion of voxel segmentations into smooth open surfaces fit for quantification was fully automated using python and pymeshlab^66^. First the individual classes in each segmentation volume were converted into unoriented point clouds, with a single point at the center of each segmented voxel. Next, the normal vectors for each point in the point cloud were estimated based on 120 neighbors with a single smoothing iteration. A surface mesh was calculated from the oriented point cloud using the screened Poisson algorithm^32^, with a reconstruction depth of 9, an interpolation weight of 4, and a minimum number of samples of 1.5. The resulting surface extended beyond the segmented region, so triangles more than 1 voxel away from the point cloud were deleted. In cases where the resulting surface was very complex, the surface was simplified with quadric edge collapse decimation to produce a surface that represented the membrane with 150,000 triangles or fewer. This process was accomplished with the ‘mrc2xyz.py’ (for point cloud generation) and ‘make_mesh.py’ scripts.

### Refinement of surface orientation and estimation of surface curvature using pycurv

Output surfaces were processed with pycurv to remove extraneous and poorly modeled regions and to refine surface normals using an area-weighted tensor voting approach^15^. This vector voting process also resulted in robust estimation of curvature metrics. Because the open surfaces generated from membrane segmentations do not have a meaningful inside or outside, signed metrics of curvature are not appropriate for analysis; instead, the unsigned curvedness metric (eq. 1) provides a single positive readout that can be used for quantification. Pycurv output triangle graph files containing the refined normal angles and curvature metrics. which enabled efficient computation of additional morphometric features, and the pycurv library was used as a base for the remaining quantifications in the manuscript. Pycurv calculations were done in the ‘calculate_curvature_args.py’ script, after file conversion using the ‘mask_and_convert_ply.py’ script.

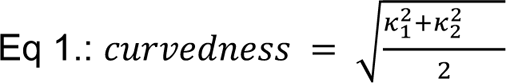

### Calculation of distances between individual surfaces

For calculations of distances between respective surface meshes, the minimum distance from each triangle on one surface to the nearest triangle on the other surface was calculated. For surfaces with small numbers of triangles, this was accomplished using a distance matrix, while calculations on large surfaces, where the distance matrix would take up more than 32GB of system memory, were accomplished using a KD-tree. Both the distance and ID of the nearest triangle on the neighboring surface are recorded in the triangle to enable further analysis. This analysis is accomplished as part of the ‘surface_to_surface.py’ script.

### Automated subclassification of the IMM

In order to understand how different components of the IMM vary with bulk morphology, triangles from inner membrane surfaces were classified into IBM, crista junction, or crista body based on their distance from the OMM. Triangles falling within 19 nm of the OMM are classified as IBM, those between 19 and 40 nm from the OMM as crista junction, and those more than 40 nm from the OMM are classified as crista body. The IBM distance was chosen as the smallest integer value above the peak OMM-IMM distance for all tomograms, while the crista body distance was selected based on visual inspection and to ensure enough triangles were assigned to the crista junctions for robust calculations.

### Calculation of distances between components within the inner membrane surface

Because all-to-all distance measurements within the same surface would trivially yield the spacing between neighboring triangles on the graph, rather than true distances between neighboring segments of membrane, measurements of distances within individual surfaces are made based on intersections along the normal vector of each triangle, similar to approaches used previously^67^. To modify this approach for measuring distances on surfaces without canonical normal directions, the first intersecting triangle was measured both along the normal vector and its inverse, and those distances were sorted into a shorter and longer distance. For cristae and junctions, the shorter distances were interpreted as the spacing between membranes within a crista or junction, whereas the longer distance of the two was interpreted as the distance between individual cristae or junctions. Normal-vector based distance measurements are sensitive to the completeness of the segmentation, and not all triangles have a normal-vector based neighbor. Those triangles were filtered out before any further quantifications.

### Calculation of relative orientations with the growth plane and neighboring regions

The smoothed normal vectors for each triangle, as estimated by pycurv, were used to estimate the orientation of mitochondrial membranes relative to each other and to the growth plane. Angles considered ranged only from 0° to 90° in the absence of canonical normal angle directions. For comparisons to the growth plane, the relative angle was calculated as follows (eq. 2):

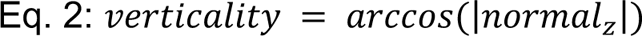

Comparisons of orientations between membranes was made by comparison of the pycurv-smoothed normal vector between each triangle on a surface to the normal vector its nearest neighbor as determined previously. The angle between these two vectors was calculated as follows (eq. 3):

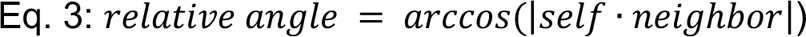

Both orientation calculations are determined in the ‘surface_to_surface.py’ script.

### Visualization and aggregate quantification of data and assessment of statistical significance

The per-triangle measurements are stored within the pycurv triangle graph file as well as being output both as surface meshes using the visualization toolkit ‘vtp’^68^ format as well as in csv files using existing pycurv functions. Visualizations were generated using the vtp files in paraview^69^. Aggregate histograms and empirical continuous distribution plots were plotted in matplotlib after aggregating all triangles across all tomograms within a condition and morphology group and weighting each triangle by its area.^70^ Violin plots were generated by generating triangle-area weighted histograms for each tomogram with 100 bins and identifying the value of the most populated bin, followed by aggregation of peak values. Statistics were similarly calculated using Mann-Whitney U tests^71^ on the peak values calculated for each tomogram to identify statistically significant differences in peak positions. For quantifying differences in distribution shape, a Mann-Whitney U test was applied to the standard deviations of each tomogram, rather than the histogram peak.

### Code Availability

All scripts used for surface reconstruction, ultrastructure quantifications, and plotting and statistical significance testing are available at https://github.com/grotjahnlab/surface_morphometrics. Scripts are licensed under the GNU Public License version 3 (GPLv3).

## REFERENCES

1. Turk, M. & Baumeister, W. The promise and the challenges of cryo-electron tomography. FEBS Lett. 594, 3243–3261 (2020).

2. Buckley, G. et al. Automated cryo-lamella preparation for high-throughput in-situ structural biology. J. Struct. Biol. 107488 (2020).

3. Schorb, M., Haberbosch, I., Hagen, W. J. H., Schwab, Y. & Mastronarde, D. N. Software tools for automated transmission electron microscopy. Nat. Methods 16, 471–477 (2019).

4. Tegunov, D., Xue, L., Dienemann, C., Cramer, P. & Mahamid, J. Multi-particle cryo-EM refinement with M visualizes ribosome-antibiotic complex at 3.5 Å in cells. Nature Methods vol. 18 186–193 (2021).

5. Himes, B. A. & Zhang, P. emClarity: software for high-resolution cryo-electron tomography and subtomogram averaging. Nat. Methods 15, 955–961 (2018).

6. Chen, M. et al. A complete data processing workflow for cryo-ET and subtomogram averaging. Nat. Methods 16, 1161–1168 (2019).

7. Jiang, Y.-F. et al. Electron tomographic analysis reveals ultrastructural features of mitochondrial cristae architecture which reflect energetic state and aging. Sci. Rep. 7, 45474 (2017).

8. Schwarz, D. S. & Blower, M. D. The endoplasmic reticulum: structure, function and response to cellular signaling. Cell. Mol. Life Sci. 73, 79–94 (2016).

9. Khanna, K., Lopez-Garrido, J., Sugie, J., Pogliano, K. & Villa, E. Asymmetric localization of the cell division machinery during Bacillus subtilis sporulation. Elife 10, (2021).

10. Ader, N. R. et al. Molecular and topological reorganizations in mitochondrial architecture interplay during Bax-mediated steps of apoptosis. Elife 8, (2019).

11. Tran, N.-H. et al. The stress-sensing domain of activated IRE1α forms helical filaments in narrow ER membrane tubes. Science 374, 52–57 (2021).

12. Mageswaran, S. K., Yang, W. Y., Chakrabarty, Y., Oikonomou, C. M. & Jensen, G. J. A cryo-electron tomography workflow reveals protrusion-mediated shedding on injured plasma membrane. Sci Adv 7, (2021).

13. Martinez-Sanchez, A., Garcia, I., Asano, S., Lucic, V. & Fernandez, J.-J. Robust membrane detection based on tensor voting for electron tomography. J. Struct. Biol. 186, 49–61 (2014).

14. Chen, M. et al. Convolutional neural networks for automated annotation of cellular cryo-electron tomograms. Nat. Methods 14, 983–985 (2017).

15. Salfer, M., Collado, J. F., Baumeister, W., Fernández-Busnadiego, R. & Martínez-Sánchez, A. Reliable estimation of membrane curvature for cryo-electron tomography. PLoS Comput. Biol. 16, e1007962 (2020).

16. Hoppe, H., DeRose, T., Duchamp, T., McDonald, J. & Stuetzle, W. Surface reconstruction from unorganized points. in Proceedings of the 19th annual conference on Computer graphics and interactive techniques 71–78 (Association for Computing Machinery, 1992).

17. Lučič, V., Rigort, A. & Baumeister, W. Cryo-electron tomography: the challenge of doing structural biology in situ. J. Cell Biol. 202, 407–419 (2013).

18. Schirrmacher, V. Mitochondria at Work: New Insights into Regulation and Dysregulation of Cellular Energy Supply and Metabolism. Biomedicines 8, (2020).

19. West, A. P. & Shadel, G. S. Mitochondrial DNA in innate immune responses and inflammatory pathology. Nat. Rev. Immunol. 17, 363–375 (2017).

20. Wai, T. & Langer, T. Mitochondrial Dynamics and Metabolic Regulation. Trends Endocrinol. Metab. 27, 105–117 (2016).

21. Chan, D. C. Mitochondrial Dynamics and Its Involvement in Disease. Annu. Rev. Pathol. 15, 235–259 (2020).

22. Sabouny, R. & Shutt, T. E. Reciprocal Regulation of Mitochondrial Fission and Fusion. Trends Biochem. Sci. 45, 564–577 (2020).

23. Rambold, A. S., Kostelecky, B., Elia, N. & Lippincott-Schwartz, J. Tubular network formation protects mitochondria from autophagosomal degradation during nutrient starvation. Proc. Natl. Acad. Sci. U. S. A. 108, 10190–10195 (2011).

24. Nielsen, J. et al. Plasticity in mitochondrial cristae density allows metabolic capacity modulation in human skeletal muscle. J. Physiol. 595, 2839–2847 (2017).

25. Colina-Tenorio, L., Horten, P., Pfanner, N. & Rampelt, H. Shaping the mitochondrial inner membrane in health and disease. J. Intern. Med. 287, 645–664 (2020).

26. Siegmund, S. E., et al. Three-Dimensional Analysis of Mitochondrial Crista Ultrastructure in a Patient with Leigh Syndrome by In Situ Cryoelectron Tomography. iScience 6, 83–91 (2018).

27. Blum, T. B., Hahn, A., Meier, T., Davies, K. M. & Kühlbrandt, W. Dimers of mitochondrial ATP synthase induce membrane curvature and self-assemble into rows. Proc. Natl. Acad. Sci. U. S. A. 116, 4250–4255 (2019).

28. Harner, M. et al. The mitochondrial contact site complex, a determinant of mitochondrial architecture. EMBO J. 30, 4356–4370 (2011).

29. Hu, C. et al. OPA1 and MICOS Regulate mitochondrial crista dynamics and formation. Cell Death Dis. 11, 940 (2020).

30. Zhang, D., Lu, C., Whiteman, M., Chance, B. & Armstrong, J. S. The mitochondrial permeability transition regulates cytochrome c release for apoptosis during endoplasmic reticulum stress by remodeling the cristae junction. J. Biol. Chem. 283, 3476–3486 (2008).

31. Frezza, C. et al. OPA1 controls apoptotic cristae remodeling independently from mitochondrial fusion. Cell 126, 177–189 (2006).

32. Kazhdan, M. & Hoppe, H. Screened poisson surface reconstruction. ACM Trans. Graph. 32, 1–13 (2013).

33. Lebeau, J. et al. The PERK Arm of the Unfolded Protein Response Regulates Mitochondrial Morphology during Acute Endoplasmic Reticulum Stress. Cell Rep. 22, 2827–2836 (2018).

34. Kremer, J. R., Mastronarde, D. N. & McIntosh, J. R. Computer visualization of three-dimensional image data using IMOD. J. Struct. Biol. 116, 71–76 (1996).

35. Navarro, P. P., Stahlberg, H. & Castaño-Díez, D. Protocols for Subtomogram Averaging of Membrane Proteins in the Dynamo Software Package. Front Mol Biosci 5, 82 (2018).

36. Bohnert, M. et al. Role of mitochondrial inner membrane organizing system in protein biogenesis of the mitochondrial outer membrane. Mol. Biol. Cell 23, 3948–3956 (2012).

37. Friedman, J. R. et al. ER tubules mark sites of mitochondrial division. Science 334, 358–362 (2011).

38. Bosc, C. et al. Autophagy regulates fatty acid availability for oxidative phosphorylation through mitochondria-endoplasmic reticulum contact sites. Nat. Commun. 11, 4056 (2020).

39. Callegari, S. et al. A MICOS-TIM22 Association Promotes Carrier Import into Human Mitochondria. J. Mol. Biol. 431, 2835–2851 (2019).

40. Bauer, M. F., Hofmann, S., Neupert, W. & Brunner, M. Protein translocation into mitochondria: the role of TIM complexes. Trends Cell Biol. 10, 25–31 (2000).

41. Aaltonen, M. J. et al. MICOS and phospholipid transfer by Ups2–Mdm35 organize membrane lipid synthesis in mitochondria. J. Cell Biol. 213, 525–534 (2016).

42. Perkins, G. et al. Electron tomography of neuronal mitochondria: three-dimensional structure and organization of cristae and membrane contacts. J. Struct. Biol. 119, 260–272 (1997).

43. Wollweber, F., von der Malsburg, K. & van der Laan, M. Mitochondrial contact site and cristae organizing system: A central player in membrane shaping and crosstalk. Biochim. Biophys. Acta Mol. Cell Res. 1864, 1481–1489 (2017).

44. Latorre-Muro, P. et al. A cold-stress-inducible PERK/OGT axis controls TOM70-assisted mitochondrial protein import and cristae formation. Cell Metab. 33, 598–614.e7 (2021).

45. Koning, R. I., Koster, A. J. & Sharp, T. H. Advances in cryo-electron tomography for biology and medicine. Ann. Anat. 217, 82–96 (2018).

46. Imhof, S. et al. Cryo electron tomography with volta phase plate reveals novel structural foundations of the 96-nm axonemal repeat in the pathogen Trypanosoma brucei. Elife 8, (2019).

47. Zhou, L. et al. Subcellular structure segmentation from cryo-electron tomograms via machine learning. bioRxiv 2020.04.09.034025 (2020) doi:10.1101/2020.04.09.034025.

48. Buchholz, T.-O., Jordan, M., Pigino, G. & Jug, F. Cryo-CARE: Content-Aware Image Restoration for Cryo-Transmission Electron Microscopy Data. arXiv [cs.CV] (2018).

49. Liu, Y.-T. et al. Isotropic Reconstruction of Electron Tomograms with Deep Learning. bioRxiv 2021.07.17.452128 (2021) doi:10.1101/2021.07.17.452128.

50. Bäuerlein, F. J. B. & Baumeister, W. Towards Visual Proteomics at High Resolution. J. Mol. Biol. 433, 167187 (2021).

51. Wietrzynski, W. et al. Charting the native architecture of Chlamydomonas thylakoid membranes with single-molecule precision. Elife 9, (2020).

52. Martinez-Sanchez, A. et al. Template-free detection and classification of membrane-bound complexes in cryo-electron tomograms. Nat. Methods 17, 209–216 (2020).

53. Basanta, B., Chowdhury, S., Lander, G. C. & Grotjahn, D. A. A guided approach for subtomogram averaging of challenging macromolecular assemblies. J Struct Biol X 4, 100041 (2020).

54. Burt, A., Gaifas, L., Dendooven, T. & Gutsche, I. Tools enabling flexible approaches to high-resolution subtomogram averaging. bioRxiv 2021.01.31.428990 (2021) doi:10.1101/2021.01.31.428990.

55. Erdmann, P. S. et al. In situ cryo-electron tomography reveals gradient organization of ribosome biogenesis in intact nucleoli. Nat. Commun. 12, 5364 (2021).

56. O’Reilly, F. J. et al. In-cell architecture of an actively transcribing-translating expressome. bioRxiv 2020.02.28.970111 (2020) doi:10.1101/2020.02.28.970111.

57. Guo, Q. et al. In Situ Structure of Neuronal C9orf72 Poly-GA Aggregates Reveals Proteasome Recruitment. Cell 172, 696–705.e12 (2018).

58. Jordan, M. A., Diener, D. R., Stepanek, L. & Pigino, G. The cryo-EM structure of intraflagellar transport trains reveals how dynein is inactivated to ensure unidirectional anterograde movement in cilia. Nat. Cell Biol. 20, 1250–1255 (2018).

59. Albert, S. et al. Direct visualization of degradation microcompartments at the ER membrane. Proc. Natl. Acad. Sci. U. S. A. 117, 1069–1080 (2020).

60. Lin, J. & Nicastro, D. Asymmetric distribution and spatial switching of dynein activity generates ciliary motility. Science 360, (2018).

61. Cignoni, P., et al. Meshlab: an open-source mesh processing tool. in Eurographics Italian chapter conference vol. 2008 129–136 (Salerno, Italy, 2008).

62. Wang, D. et al. A small molecule promotes mitochondrial fusion in mammalian cells. Angew. Chem. Int. Ed Engl. 51, 9302–9305 (2012).

63. Barad, B. Automate GIS Application Time for Cryo FIB Milling. (Github).

64. Schaffer, M. et al. Cryo-focused Ion Beam Sample Preparation for Imaging Vitreous Cells by Cryo-electron Tomography. Bio Protoc 5, (2015).

65. Mastronarde, D. N. & Held, S. R. Automated tilt series alignment and tomographic reconstruction in IMOD. J. Struct. Biol. 197, 102–113 (2017).

66. Muntoni, A., jmespadero, Luaces, A., RichardScottOZ & luzpaz. cnr-isti-vclab/PyMeshLab: PyMeshLab v2021.10. (Zenodo, 2021). doi:10.5281/ZENODO.4438750.

67. Collado, J. et al. Tricalbin-Mediated Contact Sites Control ER Curvature to Maintain Plasma Membrane Integrity. Dev. Cell 51, 476–487.e7 (2019).

68. Schroeder, W., Martin, K. & Lorensen, B. The visualization toolkit, 4th edn. Kitware. New York (2006).

69. Ahrens, J., Geveci, B. & Law, C. Paraview: An end-user tool for large data visualization. The visualization handbook 717, (2005).

70. Hunter. Matplotlib: A 2D Graphics Environment. 9, 90–95 (2007).

71. Mann, H. B. & Whitney, D. R. On a Test of Whether one of Two Random Variables is Stochastically Larger than the Other. Ann. Math. Stat. 18, 50–60 (1947).

