## Supplemental Figures for "A surface morphometrics toolkit to quantify organellar membrane ultrastructure using cryo-electron tomography"

Vehicle Elongated

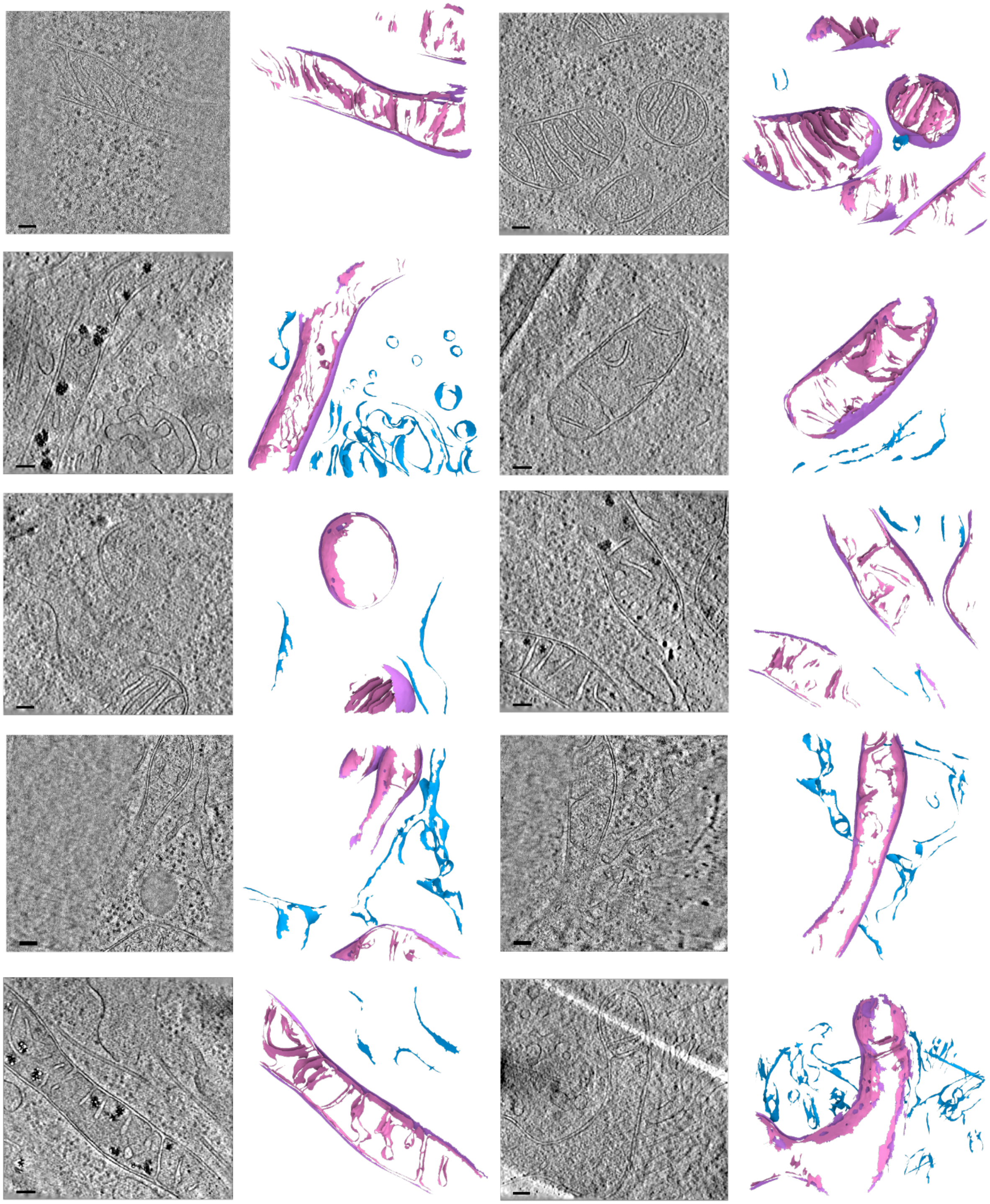

Vehicle Fragmented

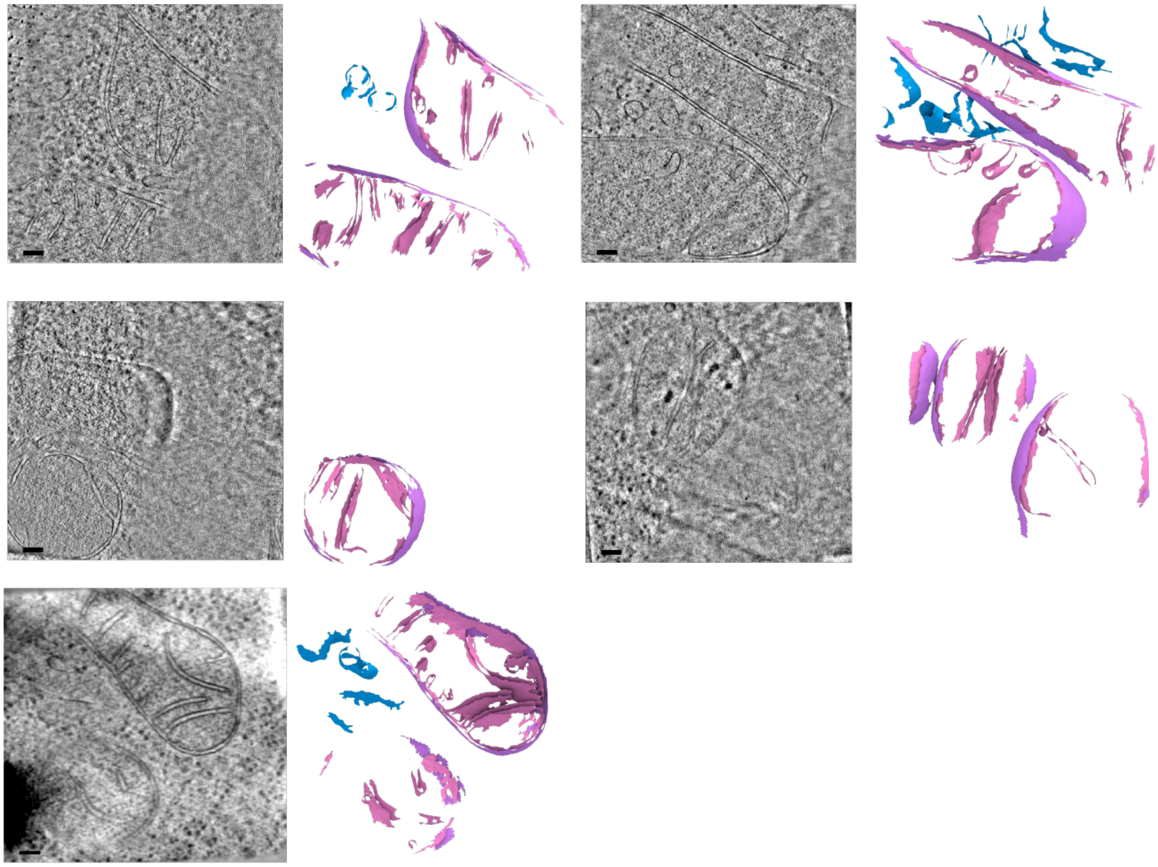

Tg Elongated

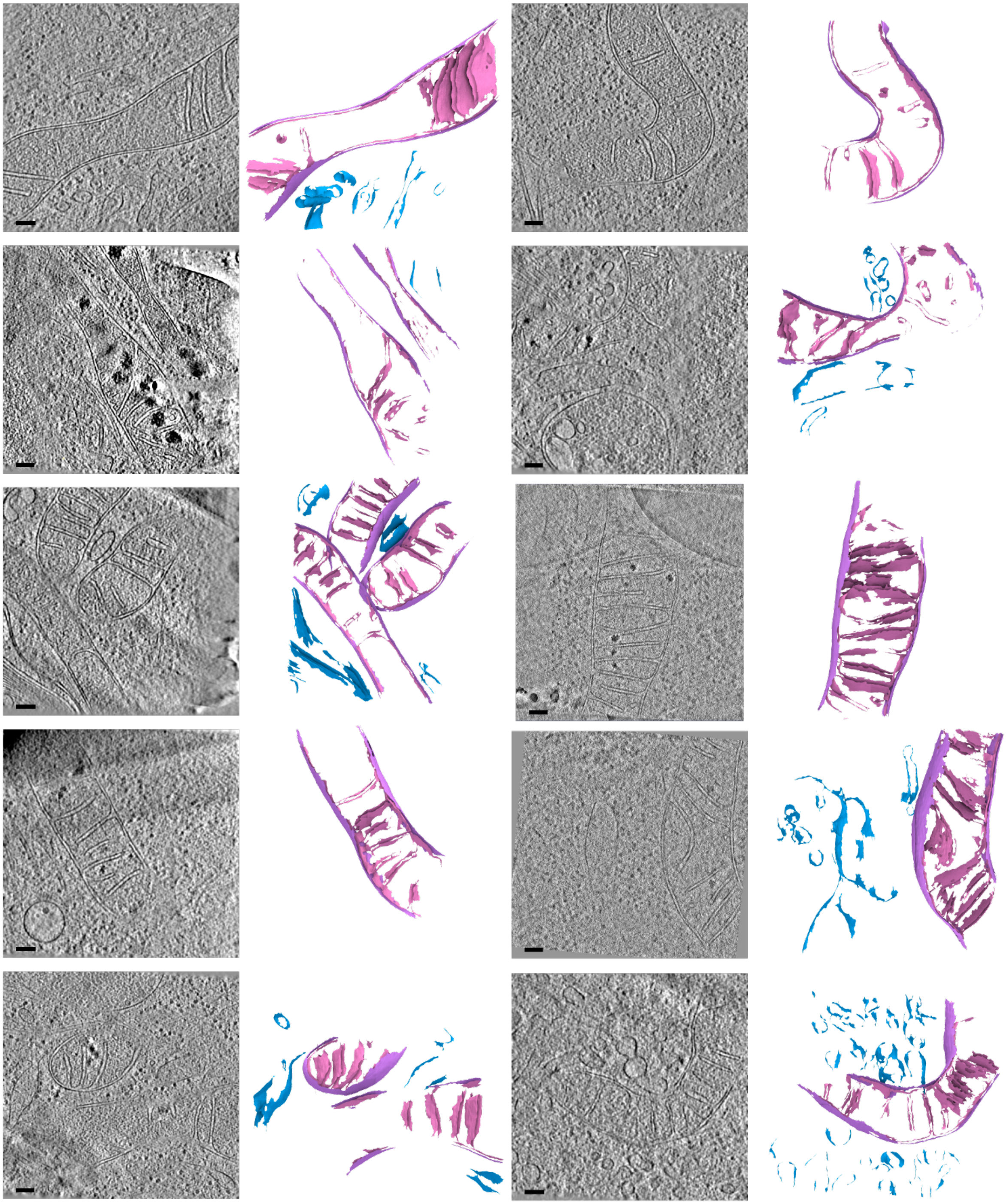

Tg Elongated (cont.)

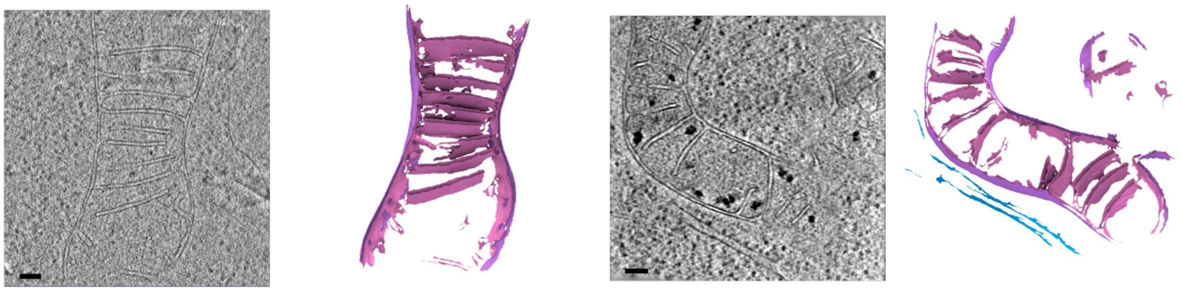

Tg Fragmented

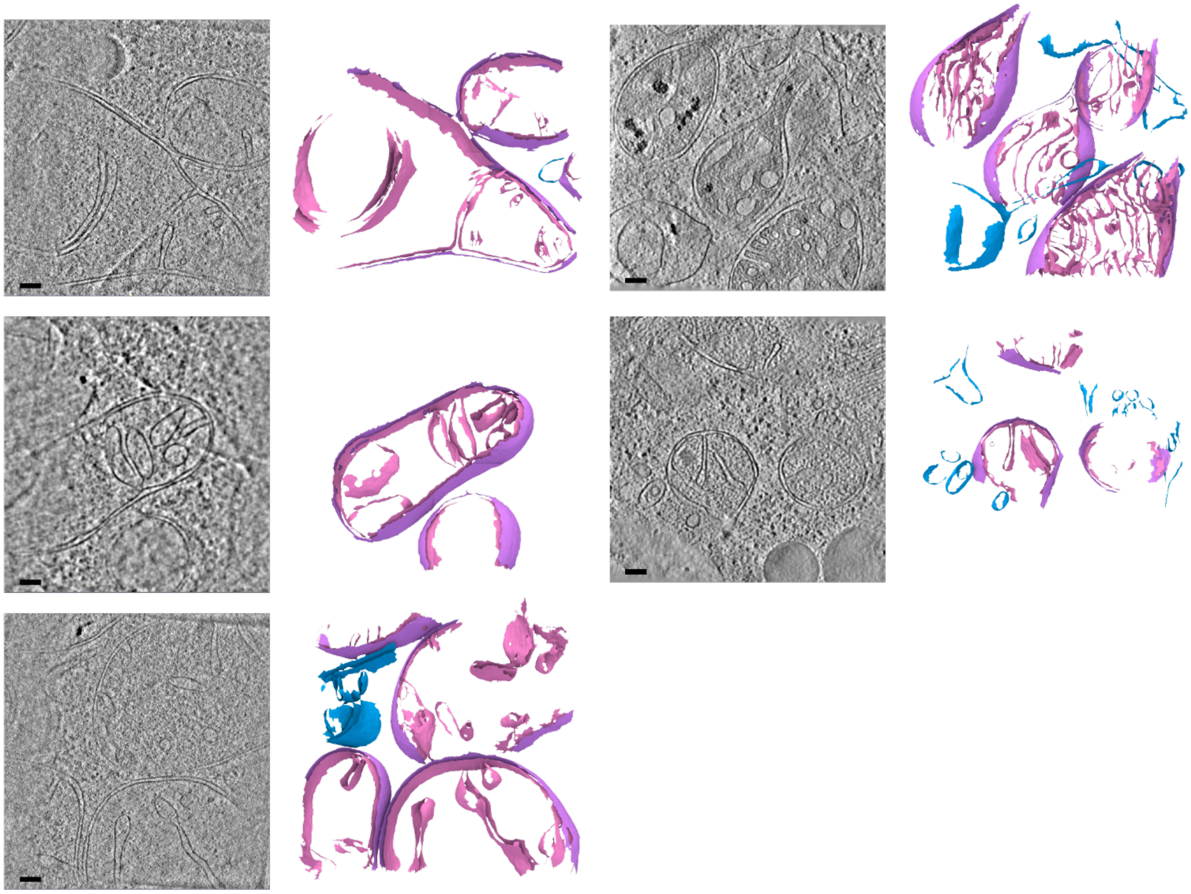

**Supplemental Figure 1. Gallery of cellular membranes used for downstream quantifications from elongated and fragmented mitochondrial networks in the presence and absence of ER stress.**

Virtual slices of tomograms (left panels) and corresponding reconstructed surface mesh models (right panels) of mitochondrial membranes (IMM, pink; OMM, purple) and ER membranes (blue) from distinct mitochondrial network morphologies (fragmented or elongated) and treatment conditions (vehicle or Tg). Meshes have been tilted backwards by 20º for visualization. Scale bar = 100 nm. (Vehicle-treated, n=15 tomograms; and Tg-treated n= 17 tomograms).

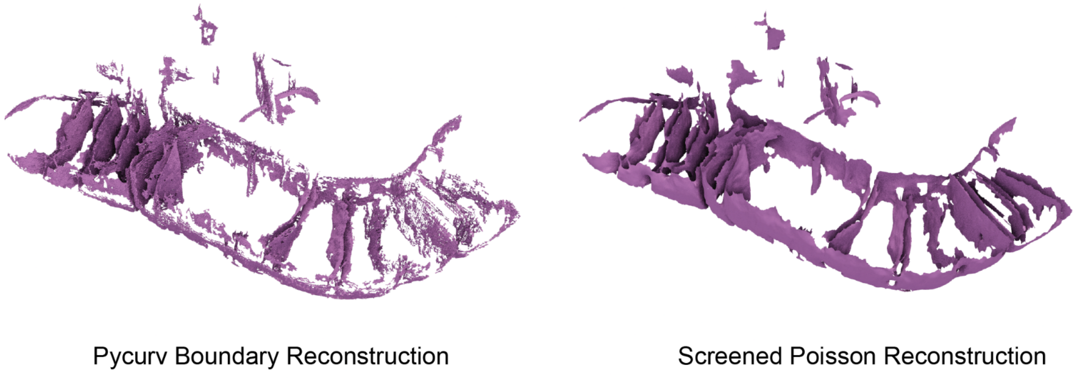


**Supplementary Figure 2. Comparison of surface reconstruction algorithms**
The same inner mitochondrial membrane segmentation was subjected to the boundary segmentation surface reconstruction algorithm used in the pycurv software[1,2] (left) as well as our new reconstruction approach based on the screened poisson algorithm[3] (right) show the improvements in smoothness and completeness enabled by the new surface reconstruction algorithm. While the left algorithm was made part of the pycurv software, it was not used for extensive analysis in the cited manuscript, in favor of algorithms using compartment segmentations.

**Supplementary Movie 1. Comparison of Voxel Segmentation vs Surface Reconstruction**
An example tomogram is shown with and without a completed voxel segmentation (green). A zoom in reveals voxel stepping artifacts that impact measurement precision, as well as small holes that result from automated segmentation. Switching to the reconstructed surface shows the smooth triangle mesh and the hole-filling properties of the surface reconstruction algorithm. Surface colors: IMM in pink, OMM in purple, ER in blue.

**References**

1. Salfer, M., Collado, J. F., Baumeister, W., Fernández-Busnadiego, R. & Martínez-Sánchez, A. Reliable estimation of membrane curvature for cryo-electron tomography. *PLoS Comput. Biol.* **16**, e1007962 (2020).
2. Hoppe, H., DeRose, T., Duchamp, T., McDonald, J. & Stuetzle, W. Surface reconstruction from unorganized points. in *Proceedings of the 19th annual conference on Computer graphics and interactive techniques* 71–78 (Association for Computing Machinery, 1992).
3. Kazhdan, M. & Hoppe, H. Screened poisson surface reconstruction. *ACM Trans. Graph.* **32**, 1–13 (2013).
